# Effects of aging on glucose and lipid metabolism in mice

**DOI:** 10.1101/2023.12.17.572088

**Authors:** Evan C. Lien, Ngoc Vu, Anna M. Westermark, Laura V. Danai, Allison N. Lau, Yetiş Gültekin, Matthew A. Kukurugya, Bryson D. Bennett, Matthew G. Vander Heiden

## Abstract

Aging is accompanied by multiple molecular changes that contribute to aging-associated pathologies, such as accumulation of cellular damage and mitochondrial dysfunction. Tissue metabolism can also change with age, in part because mitochondria are central to cellular metabolism. Moreover, the co-factor NAD^+^, which is reported to decline across multiple tissue types during aging, plays a central role in metabolic pathways such as glycolysis, the tricarboxylic acid cycle, and the oxidative synthesis of nucleotides, amino acids, and lipids. To further characterize how tissue metabolism changes with age, we intravenously infused [U-^13^C]-glucose into young and old C57BL/6J, WSB/EiJ, and Diversity Outbred mice to trace glucose fate into downstream metabolites within plasma, liver, gastrocnemius muscle, and brain tissues. We found that glucose incorporation into central carbon and amino acid metabolism was robust during healthy aging across these different strains of mice. We also observed that levels of NAD^+^, NADH, and the NAD^+^/NADH ratio were unchanged in these tissues with healthy aging. However, aging tissues, particularly brain, exhibited evidence of up-regulated fatty acid and sphingolipid metabolism reactions that regenerate NAD^+^ from NADH. Because mitochondrial respiration, a major source of NAD^+^ regeneration, is reported to decline with age, our data supports a model where NAD^+^-generating lipid metabolism reactions may buffer against changes in NAD^+^/NADH during healthy aging.

## Introduction

Aging is a time-dependent functional decline associated with increased susceptibility to pathology including diabetes, obesity, neurodegenerative diseases, cardiovascular diseases, and cancer. Several molecular hallmarks of aging have been characterized, including accumulation of cellular damage that contributes to tissue and organismal aging (López-Otín et al., 2013). One well-described hallmark is a progressive decline in mitochondrial function as organisms age with decreased mitochondrial electron transport chain function to support cellular respiration and ATP generation (Amorim et al., 2022; Green et al., 2011; López-Otín et al., 2013). Mitochondrial respiration also enables regeneration of the co-factor NAD^+^ (Luengo et al., 2021), and NAD^+^ has emerged as a critical molecule implicated in aging since its levels are reported to decline with age in many tissues in multiple organisms (McReynolds et al., 2020). Based on these observations, administration of NAD^+^ precursors such as nicotinamide riboside and nicotinamide mononucleotide is being explored as a potential therapy to extend longevity and health span (Fang et al., 2016; Mills et al., 2016).

The proposed mechanisms downstream of NAD^+^ depletion that contribute to aging are largely focused around the role of NAD^+^ as a co-substrate for enzymes such as sirtuins and poly(ADP-ribose) polymerases (Verdin, 2015). However, NAD^+^ also plays a central role in cell metabolism, serving as an essential cofactor for redox reactions involved in glycolysis, the tricarboxylic acid (TCA) cycle, and the oxidative synthesis of macromolecules such as nucleotides, amino acids, and lipids (Bao et al., 2016; Birsoy et al., 2015; Diehl et al., 2019; Li et al., 2022; Sullivan et al., 2015; Titov et al., 2016; Vander Heiden and DeBerardinis, 2017). Therefore, both aging-associated mitochondrial dysfunction and a decline in NAD^+^ levels might be expected to alter cellular metabolism in ways that may contribute to the aging process. Indeed, several studies have characterized how various metabolic processes change with age (Anderson and Weindruch, 2010; Gomes et al., 2013; Goyal et al., 2017; Hertel et al., 2016; Houtkooper et al., 2011; Laye et al., 2015; López-Otín et al., 2016; Ross et al., 2010; Walters et al., 2018).

One approach for characterizing metabolism in vivo is using intravenous infusion of isotope-labeled nutrients to trace the fate of infused nutrients into downstream metabolites within tissues by mass spectrometry (MS) (Bartman et al., 2023, 2021). This technique has been used to better understand nutrient utilization by normal tissues and tumors (Faubert et al., 2021; Hui et al., 2020, 2017). Here, we infused [U-^13^C]-glucose into healthy young and old mice to explore how tissue glucose utilization changes during aging. We examined liver, gastrocnemius muscle, and brain tissues across three different mouse strains: C57BL/6J mice, the longer-lived WSB/EiJ in-bred mouse strain (Yuan et al., 2009), and genetically heterogeneous Diversity Outbred (DO) mice. Surprisingly, we find that glucose incorporation into central carbon and amino acid metabolism is robust across these diverse strains of aging mice. Moreover, we did not measure significant declines in the levels of NAD^+^, NADH, or a change in NAD+/NADH ratio across the tissues analyzed. Rather, we observed higher levels of unsaturated fatty acids and sphingolipids in aged tissues, particularly the brain, that are generated via fatty acid desaturation and sphingolipid synthesis reactions that regenerate NAD^+^ from NADH. Given that mitochondrial respiration, which is a major source of NAD^+^ production, has been shown to decline with age in mice (Amorim et al., 2022; Green et al., 2011; López-Otín et al., 2013), our data supports a model where NAD^+^-regenerating reactions in lipid metabolism may serve as a buffer to maintain the NAD^+^/NADH ratio during healthy aging.

## Results

### Determining an optimal [U-^13^C]-glucose infusion rate to trace glucose fate in tissues in aging mice

To study how glucose fate changes in tissues in aging mice, we first determined an optimal [U-^13^C]-glucose infusion rate that would allow valid tissue metabolite labeling comparisons between young and old mice. Because older animals tend to develop insulin resistance, which impacts whole-body metabolism (Ehrhardt et al., 2019; Reynolds et al., 2019), we wanted to avoid examining changes in glucose metabolism that result from differences in insulin signaling responses between young and old mice. Therefore, we sought a glucose infusion protocol that minimally impacts whole-body metabolism and avoids effects on blood glucose and insulin levels, while still labeling sufficient metabolites in tissues to permit analysis. In a previous study evaluating the contribution of glucose to the metabolism of lung adenocarcinoma tumors in vivo, we found that a glucose infusion rate of 30 mg/kg/min allowed substantial labeling of tumor metabolites (Davidson et al., 2016). Using this rate as a starting point, we also tested rates of 15 mg/kg/min and 6 mg/kg/min (Fig. 1A). In addition, we tested two fixed infusion rates that were not adjusted for animal body weights of 0.4 mg/min and 0.2 mg/min (Fig. 1A). During a 4 h infusion in C57BL/6J mice, only the 30 mg/kg/min infusion rate increased blood glucose levels, whereas both the 30 mg/kg/min and 15 mg/kg/min infusion rates raised plasma insulin levels (Fig. 1A). When evaluating the labeling of downstream metabolites by [U-^13^C]-glucose in liver, gastrocnemius, and brain tissues, reducing the infusion rate to 6 mg/kg/min or 0.2 mg/min resulted in reduced tissue labeling of lactate and the TCA cycle metabolite succinate, whereas a rate of 0.4 mg/min led to downstream metabolite labeling that is comparable to the 30 mg/kg/min infusion rate without elevating blood glucose or insulin levels (Fig. 1A, B).

**Figure 1.**
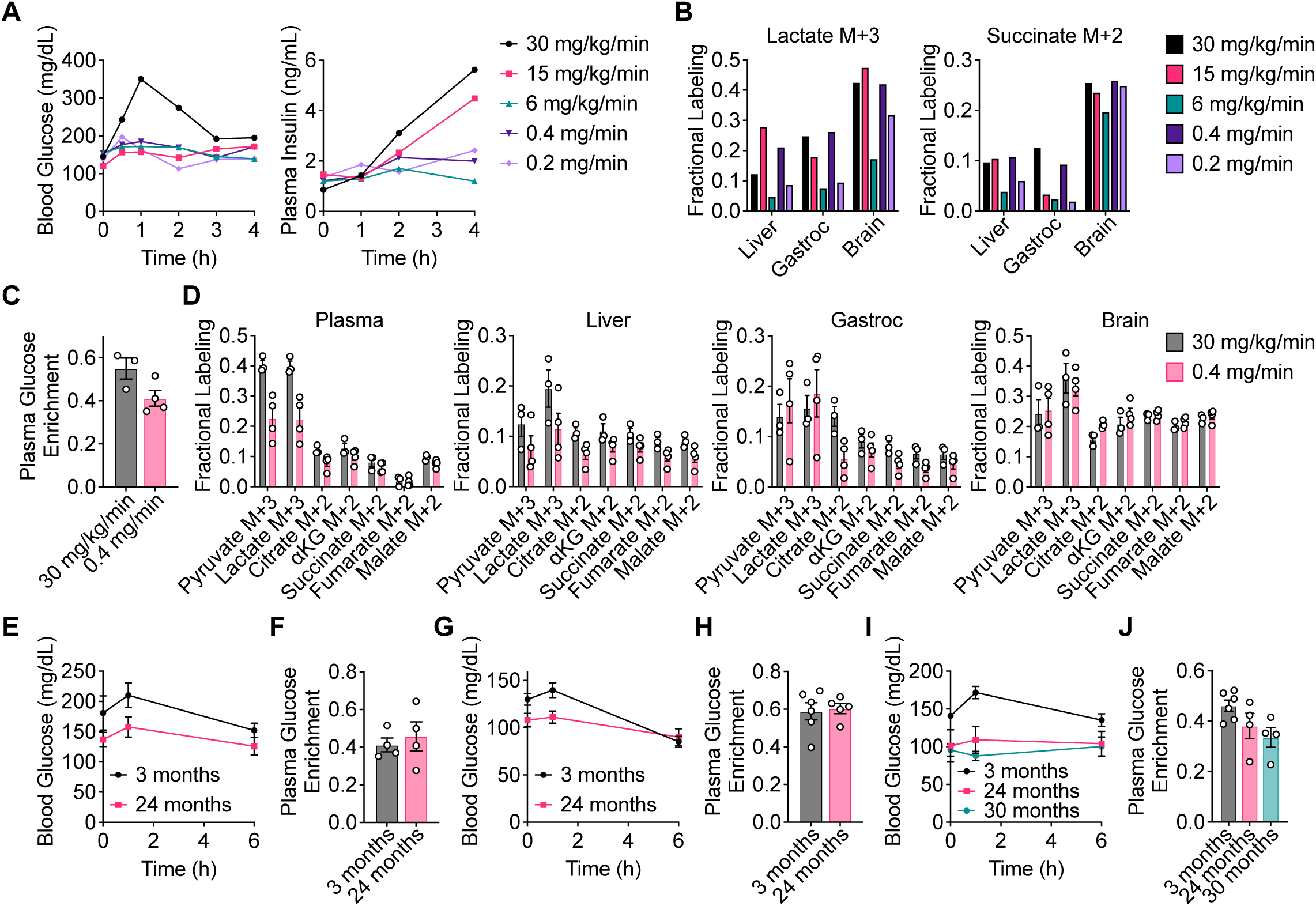
Determining an optimal [U-^13^C]-glucose intravenous infusion rate in aging mice. **A,** Serial sampling of blood glucose and plasma insulin levels over time in C57BL/6J mice infused with [U-^13^C]-glucose with the indicated infusion rates. n = 1 mouse per infusion rate. **B,** Fractional labeling of [M+3] lactate and [M+2] succinate in liver, gastrocnemius muscle (gastroc), and brain tissues from C57BL/6J mice infused with [U-^13^C]-glucose with the indicated infusion rates for 4 h. n = 1 mouse per infusion rate. **C, D,** Plasma glucose enrichment (**C**) and fractional labeling of the indicated metabolite isotopomers in plasma, liver, gastroc, and brain tissues (**D**) from C57BL/6J mice infused with [U-^13^C]-glucose at 30 mg/kg/min for 4 h (n = 3) or 0.4 mg/min for 6 h (n = 4). **E, F,** Serial sampling of blood glucose levels over time (**E**) and plasma glucose enrichment (**F**) from 3 month old (n = 4) versus 24 month old (n = 4) C57BL/6J mice infused with [U-^13^C]-glucose at 0.4 mg/min for 6 h. **G, H,** Serial sampling of blood glucose levels over time (**G**) and plasma glucose enrichment (**H**) from 3 month old (n = 6) versus 24 month old (n = 5) WSB/EiJ mice infused with [U-^13^C]-glucose at 0.4 mg/min for 6 h. **I, J,** Serial sampling of blood glucose levels over time (**I**) and plasma glucose enrichment (**J**) from 3 month old (n = 6), 24 month old (n = 4), and 30 month old (n = 4) DO mice infused with [U-^13^C]-glucose at 0.4 mg/min for 6 h. Data are presented as mean ± SEM.

We next confirmed in a larger cohort of mice that an infusion rate of 0.4 mg/min is comparable to our previously used rate of 30 mg/kg/min to allow sufficient tissue labeling for analysis. Both infusion rates resulted in substantial plasma [U-^13^C]-glucose enrichment, with 0.4 mg/min leading to only slightly lower enrichment in plasma glucose than 30 mg/kg/min (Fig. 1C). Labeling of downstream metabolites including pyruvate, lactate, and TCA cycle intermediates in plasma, liver, gastrocnemius, and brain tissues was also comparable between these infusion rates (Fig. 1D). Therefore, an infusion rate of 0.4 mg/min was determined to label metabolites to sufficient levels in tissues for analysis without significantly raising blood glucose and plasma insulin.

Since different tissues may have different nutrient uptake and exchange rates, achieving steady-state labeling is necessary for making valid comparisons between different tissues, and would allow us to evaluate whether aging alters the direct and indirect contributions of glucose to the pools of various downstream metabolites. To confirm that labeling reached steady-state in blood and tissues, we infused C57BL/6J mice with [U-^13^C]-glucose for 0.5, 2, 4, and 6 hours. Plasma [U-^13^C]-glucose enrichment reached steady-state by 6 h of infusion (Fig. S1A). Similarly, labeling of pyruvate, lactate, TCA cycle intermediates, and non-essential amino acids also reached steady-state by 6 h in plasma, liver, gastrocnemius, and brain tissues (Fig. S1B-D). Therefore, a [U-^13^C]-glucose infusion in mice at a rate of 0.4 mg/min for 6 h is able to achieve steady-state labeling of these metabolites in mouse tissues.

Finally, we assessed whether different [U-^13^C]-glucose infusion rates would impact metabolite labeling in tissues from young versus old C57BL/6J mice. In mice infused at a rate of 30 mg/kg/min, labeling of pyruvate and lactate in tissues, particularly gastrocnemius and brain, was higher in 24-month-old compared to 3-month-old mice (Fig. S2A-B). Interestingly, this difference was no longer observed in mice infused at a rate of 0.4 mg/min (Fig. 2A). These observations suggest that infusion rate differences may indeed alter tissue metabolite labeling patterns, and identifying an optimal [U-^13^C]-glucose infusion rate of 0.4 mg/min for 6 h that does not influence blood glucose and insulin levels, and reaches steady-state tissue labeling, is important for comparing tissue metabolite labeling between young and old mice.

**Figure 2.**
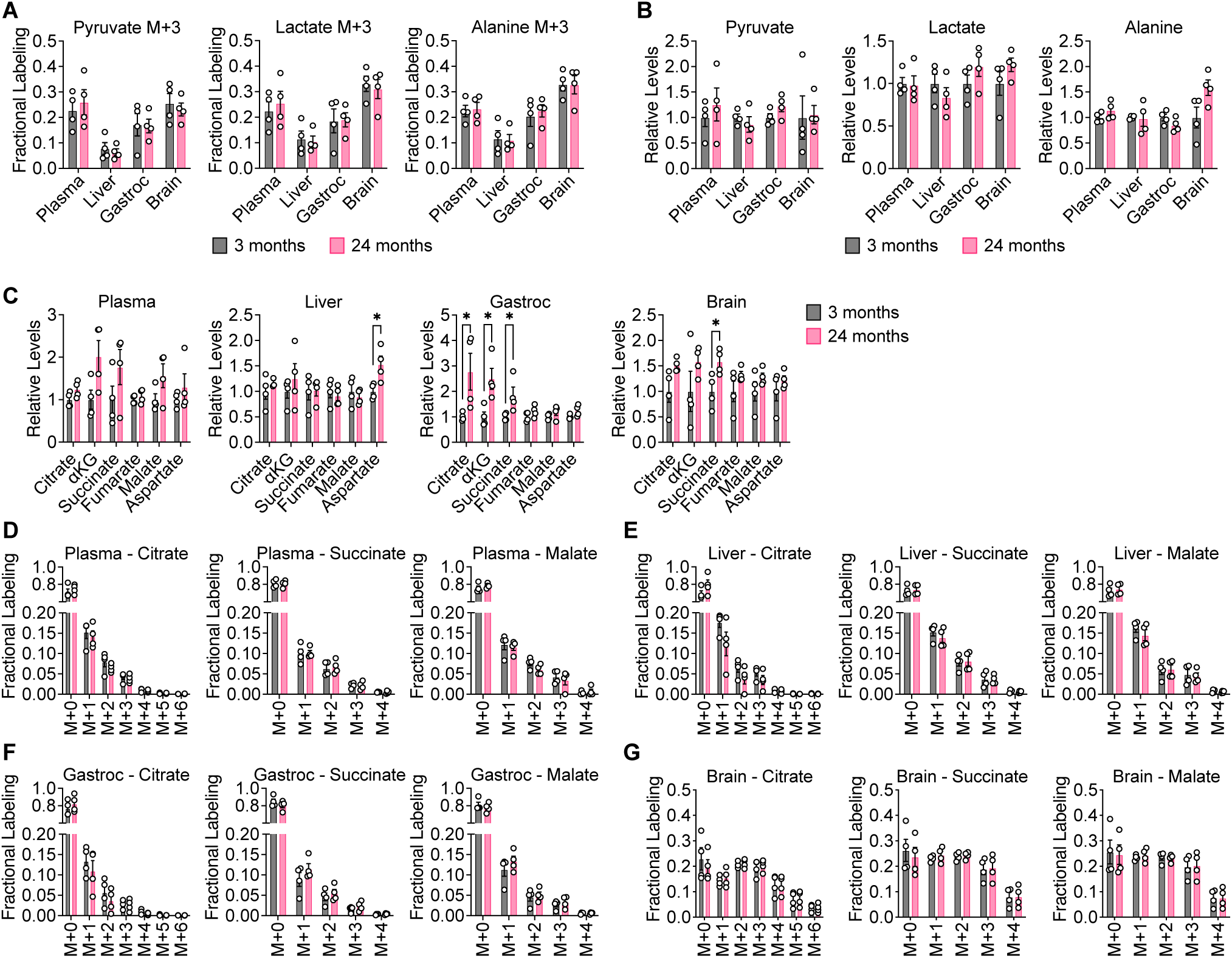
Glucose contribution to central carbon metabolism is robust in aging C57BL/6J mice. 3 month old (n = 4) versus 24 month old (n = 4) C57BL/6J mice were infused with [U-^13^C]-glucose at 0.4 mg/min for 6 h. **A, B,** [M+3] fractional labeling (**A**) and relative levels (**B**) of pyruvate, lactate, and alanine in the indicated tissues. **C,** Relative levels of TCA cycle metabolites in the indicated tissues. **D-G,** Mass isotopomer distributions of citrate, succinate, and malate in plasma (**D**), liver (**E**), gastrocnemius muscle (**F**), and brain (**G**) tissues. Data are presented as mean ± SEM. Comparisons were made using a two-tailed Student’s t-test.

### Glucose contribution to central carbon metabolism is robust in aging mice

Having established infusion parameters that enable comparison of glucose fate in young versus old mice, we conducted steady-state [U-^13^C]-glucose infusions in young and old cohorts of three distinct mouse strains. For C57BL/6J and WSB/EiJ mice, 3 month old versus 24 month old mice were compared. For diversity outbred (DO) mice, 3 month old, 24 month old, and 30 month old mice were compared. Importantly, none of the older mice exhibited overt signs of disease, allowing us to evaluate how glucose metabolism changes with healthy aging. Mice were fasted for 4 h prior to infusions, and for all mouse strains, older mice had lower fasting blood glucose at the beginning of the infusion (Fig. 1E, G, I). As expected, the infusions did not raise blood glucose levels, and for each mouse strain similar plasma [U-^13^C]-glucose enrichment was obtained between young and old mice (Fig. 1E-J).

When comparing differences in glucose labeling across animals, we first examined the contribution of glucose to the glycolytic products pyruvate and lactate in aging mice. We also considered labeling of alanine, which can be produced from pyruvate by transamination. In C57BL/6J mice, labeling of [M+3] pyruvate, lactate, and alanine was unchanged in plasma, liver, gastrocnemius, and brain tissues between young and old mice (Fig. 2A). Total levels of these metabolites also did not change in an age-dependent manner (Fig. 2B). Similar results were observed in WSB/EiJ (Fig. S3A-F) and DO mice (Fig. S3G-L). These data suggest that the contribution of glucose carbon to glycolysis in plasma, liver, gastrocnemius, and brain tissues remains robust in mice during healthy aging.

We next evaluated the contribution of glucose carbon to the TCA cycle intermediates citrate, ⍺-ketoglutarate (⍺KG), succinate, fumarate, and malate. We also considered labeling of aspartate, which can be produced from oxaloacetate by transamination. In C57BL/6J mice, total levels of these metabolites may slightly increase with age in some tissues, particularly in gastrocnemius muscle and brain (Fig. 2C). These differences, however, were not observed in WSB/EiJ and DO mice. In WSB/EiJ mice, levels of TCA cycle intermediates decreased with age in some tissues, particularly in the brain (Fig. S5A-D), whereas in DO mice, no robust age-dependent changes were observed (Fig. S6A-D). In each mouse strain, examining all mass isotopologues of TCA cycle intermediates in plasma, liver, gastrocnemius, and brain tissues revealed no differences between young and old mice in the labeling of these metabolites (Fig. 2D-G, S4A-D, S5E-H, S6E-H). These results indicate that the contribution of glucose carbon to the TCA cycle in plasma, liver, gastrocnemius, and brain tissues remains robust in mice during healthy aging.

Despite the lack of age-dependent changes, we observed several age-independent differences in tissue TCA cycle metabolite labeling. In plasma, liver, and gastrocnemius, the most abundant mass isotopologue for most TCA cycle intermediates was the [M+1] isotopologue (Fig. 2D-F, S4A-C, S5E-G, S6E-G). This in vivo labeling pattern is distinct from [U-^13^C]-glucose labeling of TCA cycle metabolites of cultured cells in vitro, in which the [M+2] and [M+3] isotopologues are typically most abundant (Davidson et al., 2016; Sellers et al., 2015). It is possible that these [M+1] species may arise from either TCA cycling or carboxylation of TCA cycle metabolites from [U-^13^C]-glucose-derived CO_2_ (Duan et al., 2022; Hensley et al., 2016). High [M+1] labeling of TCA cycle intermediates from in vivo [U-^13^C]-glucose tracing has also been observed in previous studies (Davidson et al., 2016; Duan et al., 2022; Hensley et al., 2016). In contrast, labeling of TCA cycle metabolites in brain tissues was unique and more closely mirrored in vitro labeling of cultured cells with substantial [M+2] and [M+3] isotopologues (Fig 2G, S4D, S5H, S6H). Moreover, a greater total fraction of TCA cycle metabolites in the brain were labeled by [U-^13^C]-glucose. This observation is consistent with the brain being a more glucose-avid tissue (Hui et al., 2017).

### Glucose contribution to amino acid metabolism is robust in aging mice

We next examined the contribution of glucose carbon to the non-essential amino acids asparagine, glutamine, glutamate, proline, serine, and glycine. In C57BL/6J mice, total levels of these metabolites did not robustly change with age, with some slight increases in liver and brain tissues of aged mice (Fig. 3A). Similarly, consistent age-dependent differences were not observed in WSB/EiJ and DO mice. Decreases in total levels of some amino acids were observed with age in some WSB/EiJ tissues, particularly in the gastrocnemius muscle and the brain (Fig. S7A), whereas no robust age-dependent changes were observed in DO mouse tissues (Fig. S8A). In all mouse strains, labeling of these non-essential amino acids in plasma, liver, gastrocnemius, and brain tissues revealed no differences between young and old mice (Fig. 3B-G, S7B-G, S8B-G). These results indicate that the contribution of glucose carbon to non-essential amino acid metabolism is also robust in mice during healthy aging. Again, similar to labeling of the TCA cycle, tissue-specific differences were observed with non-essential amino acid labeling. Compared to plasma, liver, and gastrocnemius tissues, brain tissues exhibited a greater contribution of glucose carbon to non-essential amino acids in all mouse strains (Fig. 3B-G, S7B-G, S8B-G), which again may be consistent with the brain being a more glucose-avid tissue (Hui et al., 2017).

**Figure 3.**
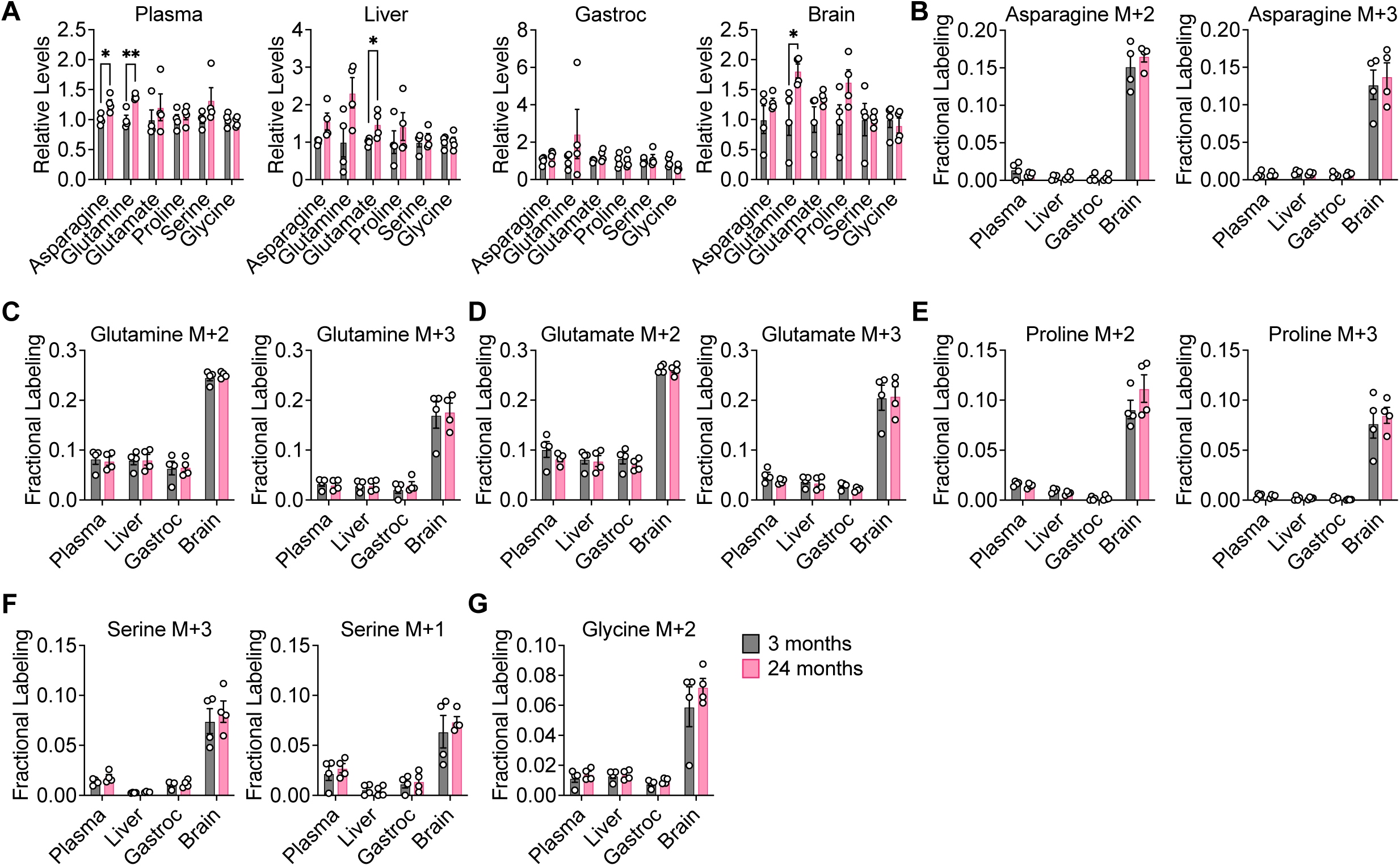
Glucose contribution to amino acid metabolism is robust in aging C57BL/6J mice. 3 month old (n = 4) versus 24 month old (n = 4) C57BL/6J mice were infused with [U-^13^C]-glucose at 0.4 mg/min for 6 h. **A,** Relative levels of the indicated amino acids in plasma, liver, gastrocnemius muscle, and brain tissues. **B-G,** Fractional labeling of [M+2] and [M+3] asparagine (**B**), [M+2] and [M+3] glutamine (**C**), [M+2] and [M+3] glutamate (**D**), [M+2] and [M+3] proline (**E**), [M+3] and [M+1] serine (**F**), and [M+2] glycine (**G**) in the indicated tissues. Data are presented as mean ± SEM. Comparisons were made using a two-tailed Student’s t-test.

### Profiling of polar metabolites does not suggest major age-dependent changes in metabolite levels

Taken together, the above data suggests that glucose contribution to central carbon and amino acid metabolism is robust across healthy aging in mice and is consistent across mouse strains, including genetically diverse DO mice. Given no major age-dependent differences in tissue metabolite labeling by [U-^13^C]-glucose, we also asked whether broader polar metabolite profiling by LC-MS would reveal any age-dependent changes in tissue metabolite levels. We compared uninfused liver, gastrocnemius, and brain tissues from 3-, 12-, and 24-month-old C57BL/6J mice. Unsupervised hierarchical clustering of metabolomics data derived from analysis of these tissues did not result in clustering of the different tissues by age (Fig. S9A-C), and principal component analyses further showed that the metabolomic profiles of liver, gastrocnemius, and brain tissues do not suggest major overall age-dependent changes (Fig. S9D-F).

### Levels of NAD^+^, NADH, and the NAD^+^/NADH ratio are stable in mouse tissues with healthy aging

Our data suggests that despite the mitochondrial dysfunction that occurs during aging (Amorim et al., 2022; Green et al., 2011; López-Otín et al., 2013), the contribution of glucose carbon, either directly or indirectly, to glycolysis, the TCA cycle, and non-essential amino acids remains robust with healthy aging. Since these measurements do not provide direct information about metabolic flux of these pathways, this conclusion is not necessarily incompatible with aging-associated impairment of mitochondrial dysfunction. Mitochondrial respiration is a major pathway for cells to regenerate NAD^+^, and impairing this process might be expected to lower the NAD^+^/NADH ratio in cells, which would be consistent with reports that NAD^+^ declines with age in multiple organisms (McReynolds et al., 2020). Therefore, we evaluated the levels of NAD^+^, NADH, and the NAD^+^/NADH ratio in aging mouse tissues. To our surprise, we found that in all three strains of mice, the NAD^+^/NADH ratio was unchanged with age in liver, gastrocnemius, and brain tissues, with the exception of NAD^+^/NADH being slightly increased in liver from C57BL/6J mice (Fig. 4A-C). Similarly, levels of NAD^+^ and NADH did not significantly change with age in all three mouse strains, with the exception of NAD^+^ being slightly increased in liver from C57BL/6J mice (Fig. 4D-F). Impaired mitochondrial respiration typically results in increased fermentation of pyruvate to lactate, which serves as an alternative route to regenerate NAD^+^. However, since the contribution of glucose carbon to [M+3] lactate in aging tissues was unchanged (Fig. 2A, S3B, S3H), we speculated that other metabolic pathways that regenerate NAD^+^ from NADH may be up-regulated during healthy aging in mice to maintain the NAD^+^/NADH ratio and compensate for any decrease in mitochondrial function.

**Figure 4.**
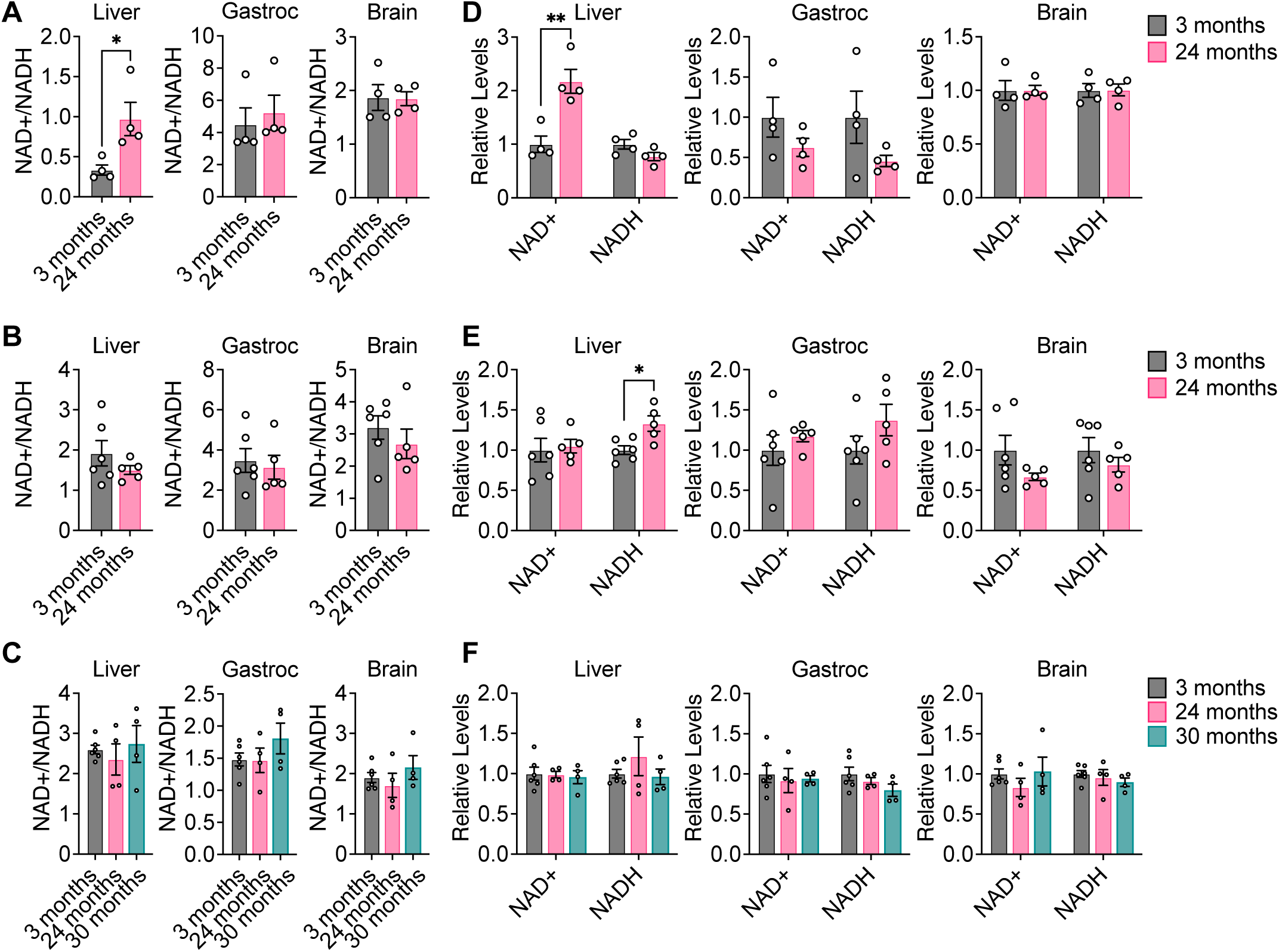
NAD^+^, NADH, and NAD^+^/NADH do not significantly change with healthy aging in mouse tissues. **A-C,** NAD^+^/NADH ratios in liver, gastrocnemius muscle, and brain tissues from young versus old C57BL/6J (**A**), WSB/EiJ (**B**), and DO (**C**) mice. **D-F,** Relative levels of NAD^+^ and NADH in in liver, gastrocnemius muscle, and brain tissues from young versus old C57BL/6J (**D**), WSB/EiJ (**E**), and DO (**F**) mice. C57BL/6J: 3 months n = 4, 24 months n = 4. WSB/EiJ: 3 months n = 6, 24 months n = 5. DO: 3 months n = 6, 24 months n = 4, 30 months n = 4. Data are presented as mean ± SEM. Comparisons were made using a two-tailed Student’s t-test.

### Fatty acid desaturation increases in aging mouse tissues

Increased membrane fatty acid desaturation in different species has been associated with decreased lifespan (Hulbert et al., 2007), and within the same species, membrane fatty acid desaturation increases with age in various tissues (Naudí et al., 2013). Moreover, caloric restriction, which robustly extends lifespan across multiple organisms, decreases membrane fatty acid desaturation in various mouse tissues (Jové et al., 2014; Lien et al., 2021; Miller et al., 2017). The production of unsaturated fatty acids is mediated by a family of fatty acid desaturase enzymes, including stearoyl-CoA desaturase (SCD), fatty acid desaturase 1 (FADS1), and fatty acid desaturase 2 (FADS2), that introduce a double bond into various fatty acid substrates (Fig. 5A). SCD produces the monounsaturated fatty acids 18:1(n-9) and 16:1(n-7) from 18:0 and 16:0, respectively. FADS1 produces polyunsaturated fatty acids, such as 20:4(n-6) from 20:3(n-6). FADS2 also primarily synthesizes polyunsaturated fatty acids, including 18:3(n-6) from 18:2(n-6), and has also been described to produce the monounsaturated fatty acid 16:1(n-10) from 16:0 (Vriens et al., 2019). Notably, these fatty acid desaturases utilize NADH as a cofactor, regenerating NAD+ when introducing a double bond into fatty acids (Fig. 5A). Upregulated production of unsaturated fatty acids may therefore maintain the NAD+/NADH ratio in tissues (Kim et al., 2019).

**Figure 5.**
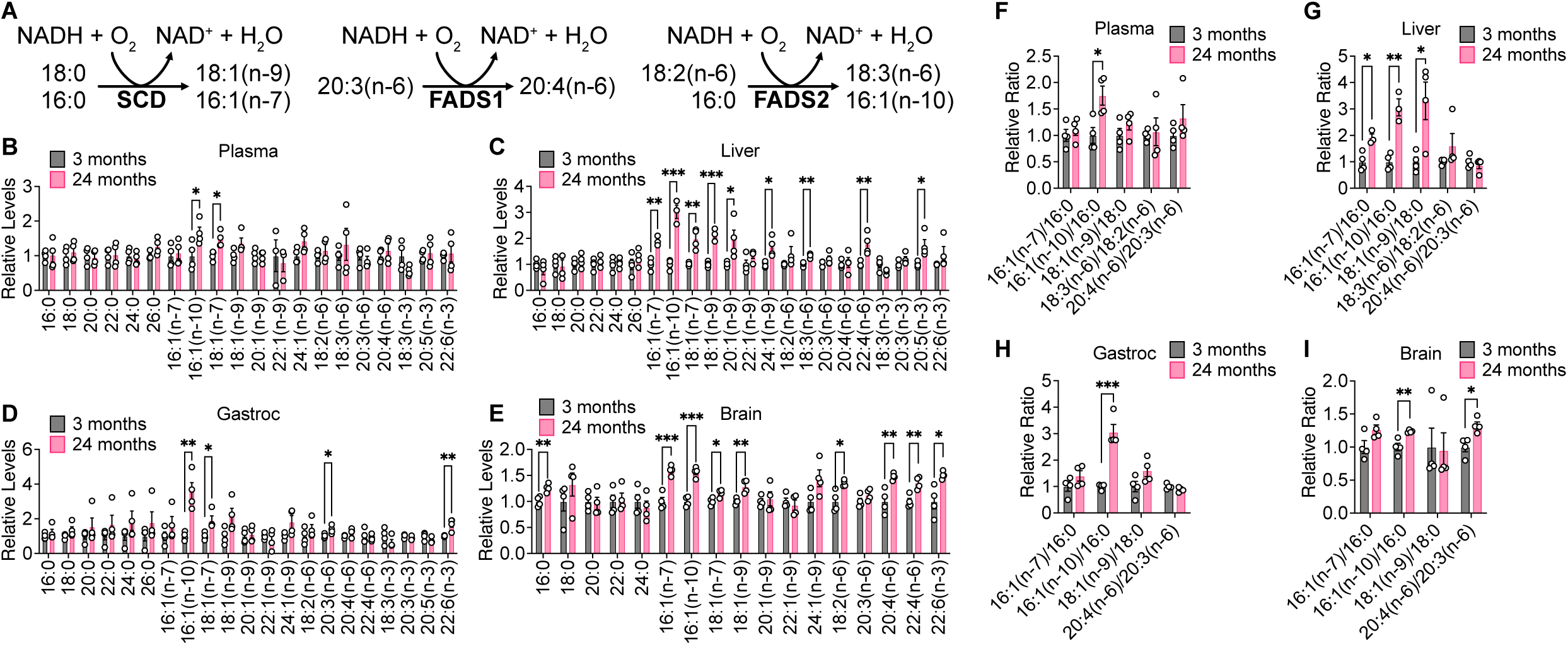
Fatty acid desaturation increases in aging C57BL/6J tissues. **A,** Schematic of fatty acid desaturation reactions by SCD, FADS1, and FADS2. **B-E,** Relative levels of the indicated fatty acids in plasma (**B**), liver (**C**), gastrocnemius muscle (**D**), and brain (**E**) tissues from young versus old C57BL/6J mice. **F-I,** Relative fatty acid desaturation ratios in plasma (**F**), liver (**G**), gastrocnemius muscle (**H**), and brain (**I**) tissues from young versus old C57BL/6J mice. 3 months n = 4, 24 months n = 4. Data are presented as mean ± SEM. Comparisons were made using a two-tailed Student’s t-test.

We examined the fatty acid composition of our aging mouse tissues by derivatizing tissue fatty acids to fatty acid methyl esters (FAME) for analysis by gas chromatography-mass spectrometry (GC-MS). Indeed, we found that aging C57BL/6J mouse tissues, particularly brain and liver tissues, have higher levels of unsaturated fatty acids (Fig. 5B-E). The ratios of the products of fatty acid desaturation reactions to their corresponding substrates have been used as surrogates for fatty acid desaturase activities (Lien et al., 2021; Vriens et al., 2019). Therefore, we assessed the 18:1(n-9)/18:0 and 16:1(n-7)/16:0 ratios as indicators of SCD activity, the 20:4(n-6)/20:3(n-6) ratio as a surrogate of FADS1 activity, and the 18:3(n-6)/18:2(n-6) and 16:1(n-10)/16:0 ratios as surrogates of FADS2 activity. These ratios all tend to increase with age in C57BL/6J tissues, particularly in the liver and brain (Fig. 5F-I). In WSB/EiJ mice, higher levels of several monounsaturated fatty acids were observed only in aged liver tissues, whereas plasma, gastrocnemius, and brain tissues exhibit few changes (Fig. S10A-D). In DO mice, increased levels of unsaturated fatty acids were not observed in aging tissues, and in gastrocnemius tissues many fatty acid species decreased with age (Fig. S11A-D). However, fatty acid desaturation ratios trended towards an increase in both aging WSB/EiJ and DO tissues (Fig. S10E-H, S11E-H), with significant increases observed in the brain in all mouse strains analyzed. Taken together, these data indicate that increased fatty acid desaturation ratios are one of the most significant age-dependent changes observed in our data sets collected across all three mouse strains, suggesting that fatty acid desaturation may increase with age in mice.

### Aging mouse brain tissue exhibits changes in sphingolipid metabolism

We next conducted lipidomics analyses on aging brain tissues because increases in fatty acid desaturation ratios were observed most robustly in the brain. Strikingly, we found that brain tissues from older mice in all three mouse strains have higher levels of sphingolipids, including hexosylceramides (HexCer), lactosylceramides (LacCer), gangliosides, and sulfatides (Fig. 6A-D). Upon considering the sphingolipid synthesis pathway, we noted that in a reaction analogous to the fatty acid desaturases, dihydroceramide desaturase (DEGS1) introduces a double bond into dihydroceramide to synthesize ceramide, which is coupled with the oxidation of NAD(P)H to NAD(P)+ (Fig. 6A). When palmitoyl-CoA is incorporated into sphingolipids by serine palmitoyltransferase (SPT), DEGS1 converts the d18:0 long chain base (LCB) in the sn-1 position of dihydroceramide into a d18:1 LCB in the sn-1 position of ceramide (Fig. 6A). We found in all three mouse strains that aged brain tissues contain overall lower levels of d18:0-containing sphingolipids and higher levels of d18:1-containing sphingolipids (Fig. S12A-C). By calculating the ratio of the total ion count of d18:1-containing lipids versus the total ion count of d18:0-containing lipids for each sphingolipid class, the d18:1/d18:0 ratios were found to be increased in aged brain tissues across all three mouse strains, particularly in HexCer and sulfatides (Fig. 6E-G). In addition, we noted that d18:2-containing sphingolipids can be generated if a monounsaturated fatty acid is incorporated at the sn-1 position of dihydroceramide by SPT that is then desaturated by DEGS1, and overall levels of d18:2-containing sphingolipids also were observed to be increased in aged brain tissues (Fig. S12A-C). Collectively, these data suggest that DEGS1 activity, which is a route for NAD^+^ regeneration, may be higher in aged mouse brain tissue.

**Figure 6.**
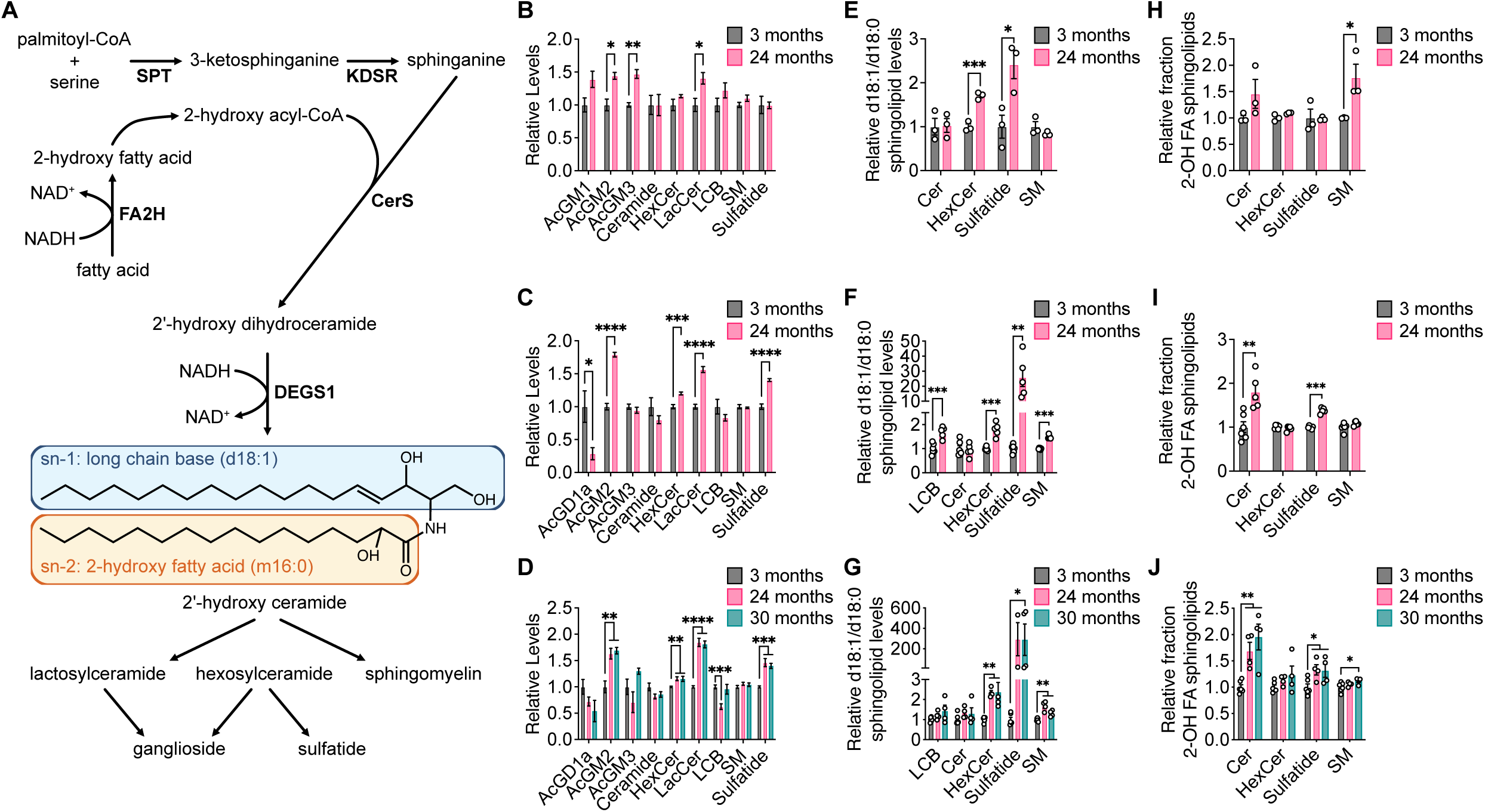
Aging brain tissues exhibit changes in sphingolipid metabolism. **A,** Schematic of sphingolipid synthesis. **B-D,** Relative levels of the indicated sphingolipid classes in brain tissues from young versus old C57BL/6J (**B**), WSB/EiJ (**C**), and DO (**D**) mice. **E-G,** Relative d18:1/d18:0 ratios of sphingolipid levels of the indicated sphingolipid classes in brain tissues from young versus old C57BL/6J (**E**), WSB/EiJ (**F**), and DO (**G**) mice. **H-J,** Relative fraction of sphingolipids containing 2-hydroxy fatty acids (2-OH FA) within each indicated sphingolipid class in brain tissues from young versus old C57BL/6J (**H**), WSB/EiJ (**I**), and DO (**J**) mice. C57BL/6J: 3 months n = 3, 24 months n = 3. WSB/EiJ: 3 months n = 6, 24 months n = 5. DO: 3 months n = 6, 24 months n = 4, 30 months n = 4. Data are presented as mean ± SEM. Comparisons were made using a two-tailed Student’s t-test. AcGM1, GM1 gangliosides; AcGM2, GM2 gangliosides; AcGM3, GM3 gangliosides; AcGD1a, GD1a gangliosides; HexCer, hexosylceramide; LacCer, lactosylceramide; LCB, long-chain base; SM, sphingomyelin.

Finally, we also detected many sphingolipid species that contain a 2-hydroxy fatty acid (2-OH FA) at the sn-2 position. 2-OH FAs are synthesized by fatty acid 2-hydroxylase (FA2H) in a reaction that is also coupled with the oxidation of NAD(P)H to NAD(P)+ (Fig. 6A). In all three mouse strains, many 2-OH FA-containing sphingolipids were found to be elevated in the brains of older mice (Fig. S13A-C). By calculating the ratio of the total ion count of 2-OH FA-containing lipids versus the total ion count of the corresponding non-hydroxylated FA-containing lipids for each sphingolipid class, we found that the relative fraction of 2-OH FA-containing sphingolipids was increased in aged brain tissues, particularly in ceramides, sulfatides, and sphingomyelins (SM) (Fig. 6H-J). These data suggest that the production of 2-OH FA by FA2H, yet another potential route for NAD^+^ regeneration, may also be higher in aged mouse brain tissue.

## Discussion

Aging is accompanied by alterations to biological processes that contribute to the accumulation of cellular damage over time (López-Otín et al., 2013). Many hallmarks of aging impinge on cellular metabolism. For example, aging-associated changes in signaling through insulin, IGF1, mTOR, and AMPK have been well-established (López-Otín et al., 2013), and these signaling pathways are major regulators of multiple metabolic pathways, including glycolysis, amino acid metabolism, and energy production (Hoxhaj and Manning, 2020). Insulin resistance tends to increase with age, leading to alterations in whole-body glucose homeostasis (Ehrhardt et al., 2019; Reynolds et al., 2019). Sirtuins, which influence longevity, also target many metabolic enzymes, including those in the electron transport chain, the TCA cycle, ketogenesis, and fatty acid oxidation (Houtkooper et al., 2012). Additionally, aging-related decline in the function of mitochondria can also impact cellular metabolism, including impaired respiration, decreased ATP generation, and increased production of reactive oxygen species (Amorim et al., 2022; Green et al., 2011; López-Otín et al., 2013). These observations led us to hypothesize that glucose utilization by tissues may change during aging. Indeed, prior work has provided evidence for aging-related changes in glucose metabolism. For example, FDG-PET imaging of human brains suggests that glucose uptake and aerobic glycolysis are impaired during aging, particularly with neurodegenerative disease (Goyal et al., 2017). On the other hand, several mouse studies have suggested that aerobic glycolysis and lactate levels increase with age in some tissues, including the brain and skeletal muscle (Gomes et al., 2013; Ross et al., 2010). Profiling of aging animals has also revealed alterations in the expression of glucose metabolism genes and the levels of metabolites involved in central carbon and amino acid metabolism in multiple tissues (Houtkooper et al., 2011; Walters et al., 2018).

The intravenous infusion of stable isotope-labeled nutrients can be a powerful tool for assessing nutrient utilization in tissues within live animals. This approach has helped expand our understanding of tumor metabolism, particularly by highlighting how in vivo tumor metabolism can be distinct from the metabolism of cultured cancer cells and how it varies across tissues (Altea-Manzano et al., 2023; Davidson et al., 2016; Elia et al., 2019; Faubert et al., 2017; Hui et al., 2017; Lau et al., 2020; Yuneva et al., 2012). This method has also been used to explore normal physiology to understand the fuel preferences of different tissues (Hui et al., 2020, 2017; Jang et al., 2019, 2018). In this study we used intravenous infusion of [U-^13^C]-glucose in young and old mice across three different mouse strains to examine how glucose utilization in normal tissues changes during aging. While minimal changes in glucose labeling were observed in tissues with age, several caveats are associated with the interpretation of these data. First, because changes to whole-body metabolic physiology occur during aging, such as increased insulin resistance, this can lead to distinct responses by young versus old mice to the infusion of exogenous glucose. We optimized a [U-^13^C]-glucose infusion rate in young and old mice that avoids elevating blood glucose and insulin levels while still leading to reasonable tissue metabolite labeling (Fig. 1), and we find that using different infusion rates can lead to distinct results in tissue metabolite labeling between young and old mice (Fig. S2). However, despite our best efforts to determine an optimal infusion rate, we cannot rule out that young and old mice may still respond differently to the glucose infusions. Another caveat is that while steady-state labeling of [U-^13^C]-glucose can provide data on relative differences in label incorporation into different metabolites, this approach does not directly measure differences in metabolic pathway fluxes. Therefore, our data argue against major shifts in fuel sources used by different tissues, but may not reflect more subtle changes in metabolic pathway fluxes. Finally, conclusions about tissue glucose utilization are complicated by inter- and intra-tissue metabolic interactions. For example, while labeling of glycolytic and TCA cycle intermediates from [U-^13^C]-glucose in the muscle can result from direct glucose uptake by muscle cells, indirect labeling can also occur through labeled intermediates that are produced by other organs and released into circulation. Similar interactions can occur between different cell types within the same tissue. Label incorporation can also be confounded by exchange flux, in which rapid substrate-product interconversion by reversible metabolic reactions can lead to labeling of downstream metabolites regardless of the net direction of those reactions (Buescher et al., 2015).

To our surprise, we find that glucose contribution to glycolysis, the TCA cycle, and amino acid metabolism in the tissues analyzed does not significantly change during healthy aging in C57BL/6J, WSB-EiJ, and DO mice (Fig. 2). The only observed difference in metabolite labeling is tissue-specific, with brain tissues exhibiting distinct labeling patterns compared to plasma, liver, and gastrocnemius that are consistent with the brain being a more glucose-avid tissue (Hui et al., 2017). While our conclusion that the utilization of glucose carbon, either directly or indirectly, as a fuel source for metabolism is robust in healthily aging mice may be unexpected, our data provides little information about metabolic pathway fluxes and thus are not inconsistent with well-documented evidence that the activities of pathways related to glucose metabolism, such as mitochondrial respiration, decline with age.

As a major pathway for NAD^+^ production, mitochondrial dysfunction may contribute to the reported age-associated decline in NAD^+^ levels (McReynolds et al., 2020). However, we find that levels of NAD^+^, NADH, and the NAD^+^/NADH ratio also do not change during healthy aging in our mouse cohorts (Fig. 4). Notably, a recent study also showed that the degree of aging-associated NAD^+^ decline can vary across tissues, with only minor or no changes to tissue NAD^+^/NADH ratios (McReynolds et al., 2021). These results may imply that other NAD^+^-regenerating metabolic reactions may compensate to maintain the appropriate NAD^+^/NADH ratio in tissues with healthy aging, and perhaps the declines observed by others may be reflective of age-related tissue dysfunction.

Our data suggests that lipid metabolism reactions may be a contributor to NAD^+^/NADH homeostasis with aging, particularly in brain tissue. In all three mouse strains, fatty acid desaturation, which converts NADH to NAD^+^ in the process of synthesizing unsaturated fatty acids, appears to be up-regulated in older tissues, particularly the brain. This conclusion is consistent with a report describing how polyunsaturated fatty acid synthesis through the fatty acid desaturases FADS1 and FADS2 are up-regulated to generate NAD^+^ in response to inhibition of mitochondrial respiration (Kim et al., 2019). Notably, previous studies have also shown that membrane fatty acid desaturation increases with age in many tissues. These highly unsaturated fatty acids are more susceptible to lipid peroxidation by reactive oxygen species, which is hypothesized to contribute to aging-related cellular damage (Naudí et al., 2013). Similarly, we also find in all three mouse strains that aging brain tissues have elevated levels of sphingolipids, with additional evidence suggesting aging-associated increases in the activities of DEGS1 and FA2H, which are coupled to the oxidation of NAD(P)H to NAD(P)^+^. These observations are consistent with prior studies observing a similar increase in the levels of sphingolipids in aging brain tissues, particularly species containing unsaturated and 2-hydroxy fatty acids (Couttas et al., 2018; Kishimoto and Radin, 1959). Ceramides in particular are lipids that can be pro-apoptotic and have been implicated in aging-related pathologies such as neurodegeneration and insulin resistance (Chaurasia and Summers, 2015).

We propose a model for understanding these aging-associated changes in fatty acid desaturation and sphingolipid metabolism in the context of NAD^+^/NADH homeostasis. In normal tissues in young animals, fully functional mitochondrial respiration enables cells to maintain a NAD^+^/NADH ratio that supports tissue function. As aging leads to progressive mitochondrial dysfunction, cells may need to shift to alternative metabolic reactions that regenerate NAD^+^ to maintain their NAD^+^/NADH ratio, including lipid metabolism reactions such as fatty acid desaturation and sphingolipid synthesis. These reactions may therefore serve as a compensatory system that buffers against mitochondria-associated declines in NAD^+^ production. Whether the availability of fatty acid and lipid precursors for desaturation and sphingolipid synthesis themselves can become limiting is not known, but evidence for ongoing membrane remodeling and lipid turnover in tissues (Wang and Tontonoz, 2019) make it likely that sustained activity of these lipid metabolism reactions is possible. It is also intriguing that while such compensatory reactions may initially serve to restore NAD^+^/NADH homeostasis, they may ultimately produce metabolites that contribute to age-associated cellular damage, such as unsaturated fatty acids that are more susceptible to peroxidation and pro-apoptotic ceramides. Importantly, we also note that the old mice analyzed in our cohorts were overtly healthy, and thus we speculate that reports in the literature of age-associated decline in NAD^+^ levels may reflect measurements from older, potentially sicker, animals in which age-related pathologies have presented that overwhelm NAD^+^/NADH buffering systems.

We acknowledge that the presence of such a NAD^+^/NADH buffering system involving lipid metabolism is difficult to test. One prediction of our model is that older animals will have already engaged alternative NAD^+^-generating reactions to maintain the NAD^+^/NADH ratio, and compared to younger animals, older animals will be more sensitive to further stresses such as mitochondrial electron transport chain inhibitors that perturb this ratio. Interestingly, the mitochondrial complex I inhibitor rotenone has been used in rat and mouse models to induce features of Parkinson’s disease (Betarbet et al., 2000), and administration of rotenone to aged mice leads to more severe Parkinson’s-like symptoms compared to young mice (Weetman et al., 2013). As further work is done to characterize how various metabolic pathways change during aging, it will be interesting to focus on reactions that are capable of regenerating NAD+ to help maintain NAD^+^/NADH homeostasis.

## Materials and Methods

### Animal studies

All experiments conducted in this study were approved by the MIT Committee on Animal Care (IACUC). Male C57BL/6J mice were obtained from The Jackson Laboratory (000664) or Calico Labs and aged in house. Male 3-month-old and 2-year-old WSB/EiJ and Diversity Outbred (DO) mice were obtained from Calico Labs. All animals were housed at ambient temperature and humidity (18-23°C, 40-60% humidity) with a 12 h light and 12 h dark cycle and co-housed with ad libitum access to water. Data was collected from distinct animals, where n represents biologically independent samples. Only mice with a body score condition of ≥2 were used for experiments. Statistical methods were not performed to pre-determine sample size.

### Glucose infusion

Infusion of [U-^13^C]-glucose (Cambridge Isotope Laboratories) was performed as previously described (Davidson et al., 2016; Lau et al., 2020). A catheter was surgically implanted into the jugular vein of animals 3-4 days prior to infusion. Mice were fasted for 4 h prior to starting infusions, which were conducted in conscious, free-moving animals at the indicated infusion rates for up to 6 h.

### Blood glucose and plasma insulin measurements

Blood glucose levels were measured using a Contour glucose meter (Ascensia Diabetes Care). Plasma insulin was measured with an ultra-sensitive mouse insulin ELISA (Crystal Chem #90080).

### Tissue and plasma polar metabolite extraction

Snap frozen tissues were ground into powder, and polar metabolites were extracted with a 5:3:5 ratio of ice-cold HPLC-grade methanol:water:chloroform containing norvaline as an internal standard. For plasma, blood collected from animals was immediately placed in EDTA tubes (Sarstedt 41.1395.105) and centrifuged to separate plasma. 10 µl of plasma was extracted with 300 µl of ice-cold methanol. Samples were vortexed for 15 min at 4°C and centrifuged at maximum speed for 10 min at 4°C. The aqueous polar metabolite fraction was dried under nitrogen gas and frozen at -80°C until analysis.

### Tissue and plasma lipid extraction for fatty acid analysis

Snap frozen tissues were ground into powder using a mortar and pestle. Tissue powder was then weighed into glass vials (Thermofisher C4010-1, C4010-60BLK). Blood collected from animals was immediately placed in EDTA tubes (Sarstedt 41.1395.105) and centrifuged to separate plasma. Lipids were extracted in 1.5 ml dichloromethane:methanol (containing 25 mg/L butylated hydroxytoluene, Millipore Sigma B1378):0.88% KCl (w/v) (8:4:3), vortexed for 15 min at 4°C, and centrifuged at maximum speed for 10 min at 4°C. The extraction buffer contained either 0.7 µg/ml tridecanoic acid or 0.7 µg/ml cis-10-heptadecenoic acid as internal standards. Lipids (organic fraction) were transferred to glass vials, dried under nitrogen gas, and immediately processed for analysis.

### Gas chromatography-mass spectrometry (GC-MS) analysis of polar metabolites

Polar metabolites were analyzed by GC-MS as described previously (Lien et al., 2021). Dried and frozen metabolite extracts were derivatized with 16 µl of MOX reagent (Thermo Fisher TS-45950) for 60 min at 37°C, followed by derivatization with 20 µl of N-tert-butyldimethylsilyl-N-methyltrifluoroacetamide with 1% tert-butyldimethylchlorosilane (Millipore Sigma 375934) for 30 min at 60°C. Derivatized samples were analyzed by GC-MS, using a DB-35MS column (Agilent Technologies 122-3832) installed in an Agilent 7890B gas chromatograph coupled to an Agilent 5997B mass spectrometer. Helium was used as the carrier gas at a constant flow rate of 1.2 mL/min. One microliter of sample was injected in split mode (1:10) at 270°C. After injection, the GC oven was held at 100°C for 1 min, increased to 105°C at 2.5°C/min, held at 105°C for 2 min, increased to 250°C at 3.5°C/min, and then ramped to 320°C at 20°C/min. The MS system operated under electron impact ionization at 70 eV, and the MS source and quadrupole were held at 230°C and 150°C, respectively. The detector was used in scanning mode with an ion range of 100-650 m/z.

Glucose was analyzed by GC-MS as described previously (Antoniewicz et al., 2011). Dried and frozen metabolite extracts were derivatized with 50 µl of 2% (w/v) hydroxylamine hydrochloride in pyridine (Millipore Sigma) for 60 min at 90°C, followed by derivatization with 100 µl of propionic anhydride (Millipore Sigma) for 30 min at 60°C. Derivatized samples were then dried under nitrogen gas and resuspended in 100 µl of ethyl acetate (Millipore Sigma) in glass GC-MS vials. Samples were analyzed by GC-MS as described above, except helium was used as the carrier gas at a constant flow rate of 1.1 mL/min, and one microliter of sample was injected in splitless mode at 250°C. After injection, the GC oven was held at 80°C for 1 min, ramped to 280°C at 20°C/min, and held at 280°C for 4 min.

Total ion counts were determined by integrating appropriate ion fragments for each metabolite using El-Maven software (Elucidata). Mass isotopologue distributions were corrected for natural abundance using IsoCorrectoR (Heinrich et al., 2018). Metabolite data was normalized to the internal standard and biofluid volumes/tissue weights.

### Gas chromatography-mass spectrometry (GC-MS) analysis of fatty acid methyl esters

Fatty acid methyl esters (FAMEs) were analyzed by GC-MS as described previously (Lien et al., 2021). Dried lipid extracts were resuspended in 100 µl of toluene in glass vials and derivatized with 200 µl of 2% sulfuric acid in methanol overnight at 50°C. After derivatization, 500 µl of 5% NaCl was added, and FAMEs were extracted twice with 500 µl of hexane. Samples from animal tissues or biofluids were cleaned up with Bond Elut LRC-Florisil columns (Agilent Technologies 12113049). Columns were pre-conditioned with 3 mL of hexane, and then the FAME extracts in hexane were added to the column. FAMEs were finally eluted twice with 1 ml of hexane:diethyl ether (95:5 v/v), dried under nitrogen gas, and resuspended in hexane for GC-MS analysis. GC-MS was conducted with a DB-FastFAME column (Agilent Technologies G3903-63011) installed in an Agilent 7890A gas chromatograph coupled to an Agilent 5975C mass spectrometer. Helium was used as the carrier gas at a constant pressure of 14 psi. One microliter of sample was injected in splitless mode at 250°C. After injection, the GC oven was held at 50°C for 0.5 min, increased to 194°C at 25°C/min, held at 194°C for 1 min, increased to 245°C at 5°C/min, and held at 245°C for 3 min. The MS system operated under electron impact ionization at 70 eV, and the MS source and quadrupole were held at 230°C and 150°C, respectively. The detector was used in scanning mode with an ion range of 104-412 m/z. Total ion counts were determined by integrating appropriate ion fragments for each FAME using El-Maven software (Elucidata). Metabolite data was background corrected using a blank sample and normalized to the internal standard and biofluid volumes/tissue weights.

### NAD+/NADH measurements

Snap frozen tissues were ground into powder using a mortar and pestle. Tissue powder was weighed and extracted in 200 µl of ice-cold lysis buffer (1% dodecyltrimethylammonium bromide in 0.2 N of NaOH diluted 1:1 with PBS) per 10 mg tissue, snap frozen in liquid nitrogen, and frozen at -80°C. NAD^+^ and NADH were measured using a protocol adapted from the manual of the NAD^+^/NADH-Glo Assay Kit (Promega, G9072), as previously described (Li et al., 2022).

### Extraction of tissue and plasma for LC-MS metabolomics and lipidomics

50 µL of plasma was used per sample, and 35 mg of homogenized tissue was used for extraction. Sample were placed in 2 mL glass vial and extracted for LC-MS-based lipidomics and metabolomics analysis using methyl-tert-butyl ether liquid-liquid extraction (MTBE-LLE) (Matyash et al., 2008). In short, 350 µL of water and 400 µL of methanol containing internal standards (100 µL of LipidSplash standards (Avanti Polar Lipids), 0.5 µg/mL Tyr-d5, 0.5 µg/mL Arg-^15^N_4_, and 125 ng/mL Benzonic acid-d5) was added to the sample, vortexed for 30 s, and incubated on ice for 15 minutes. 800 µL of MTBE was added to the mixture, vortexed for 30 s, incubated on ice for another 15 minutes, and centrifuged at 3500 RPM for 10 minutes at 4°C. Lipids partitioned in the top layer was collected into a separate vial, and the extraction process was repeated again with an additional 600 µL of MTBE. After incubating and centrifuging, the second organic layer was collected and combined in the first lipid vial. The remaining aqueous layer contains polar metabolites was transferred to a glass vial. Both extractions were dried under nitrogen at 4°C. For lipidomics analysis, samples were resuspended in 200 µL of butanol/methanol/water (2:1:1, v/v/v), and polar metabolites were resuspended in 80% acetonitrile for LC-MS analysis.

### LC-MS analysis of polar metabolites

Metabolomics samples were analyzed in both positive and negative ESI-LC-MS methods on Vanquish UPLCs coupled to Q-Exactive Plus mass spectrometers. In positive mode, metabolites were separated using a SeQuant^®^ ZIC^®^-pHILIC column (5 μm, 200 Å, 150 × 2.1 mm). Mobile phase A was 20 mM ammonium carbonate in water (pH 9.2) and mobile phase B was acetonitrile at a flow rate of 150 μL/min and the gradient was t = −6, 80% B; t = 0, 80% B; t = 2.5, 73% B; t = 5, 65% B, t = 7.5, 57% B; t = 10, 50% B; t = 15, 35% B; t = 20; 20% B; t = 22, 15% B; t = 22.5, 80% B; t = 24; 80% B. Data was acquired using data-dependent acquisition (DDA) mode with the following parameters: resolution = 70,000, AGC target = 3.00 × 10^6^, maximum IT (ms) = 100, scan range = 70–1050. The MS2 parameters were as follows: resolution = 17,500, AGC target = 1.00 × 10^5^, maximum IT (ms) = 50, loop count = 6, isolation window (*m*/*z*) = 1, (N)CE = 20, 40, 80; underfill ratio = 1.00%, Apex trigger(s) = 3–10, dynamic exclusion(s) = 25.

In negative mode, metabolites were separated using a reverse phase ion-pairing chromatographic method using an Agilent Extend C18 RRHD column (1.8 μm, 80 Å, 2.1 × 150 mm). Mobile phase A was 10 mM tributylamine, 15 mM acetic acid in 97:3 water:methanol pH 4.95; mobile phase B was methanol at a flow rate of 200 μL/min. The gradient was t = −4, 0% B; t = 0, 0% B; t = 5; 20% B; t = 7.5, 20% B; t = 13, 55% B; t = 15, 95% B; t = 18.5, 95% B; t = 19, 0% B; t = 22, 0% B. Data was acquired in data-dependent acquisition (DDA) mode with the following parameters: resolution = 70,000, AGC target = 1.00 × 10^6^, maximum IT (ms) = 100, scan range = 70–1050. The MS/MS parameters were as follows: resolution = 17,500, AGC target = 1.00 × 10^5^, maximum IT (ms) = 50, loop count = 6, isolation window (*m*/*z*) = 1, (N)CE = 20, 50, 100; underfill ratio = 1.00%, Apex trigger(s) = 3–12, dynamic exclusion(s) = 20.

### LC-MS analysis of lipids

Lipidomics samples were analyzed in both positive and negative ion mode using the same LC-MS method consisting of a Vanquish UPLC coupled to a Q-Exactive Plus mass spectrometer. Lipids were separated using a Thermo Scientific Accucore C30 column (2.6 μm, 150 Å, 2.1 × 250 mm) at a flow rate of 200 μL/min. For the the LC method, mobile phase A was 20 mM ammonium formate in 60:40 acetonitrile:water + 0.25 μM medronic acid, and mobile phase B was 20 mM ammonium formate in 90:10 isopropanol:acetonitrile + 0.25 μM medronic acid. The gradient was t = −7, 30% B, t = 0, 30% B, t = 7, 43% B, t = 12, 65% B, t = 30, 70% B, t = 31, 88% B, t = 51, 95% B, t = 53, 100% B, t = 55, 100% B, t = 55.1, 30% B, t = 60, 30% B. The mass spectrometer settings were as follows: data-dependent acquisition (DDA) was performed with the following parameters: resolution = 140,000, AGC target = 3.00 × 10^6^, maximum IT (ms) = 100, scan range = 200–2000. The MS2 parameters were as follows: resolution = 17,500, AGC target = 3.00 × 10^6^, maximum IT (ms) = 150, loop count = 8, isolation window (*m*/*z*) = 1, (N)CE = 20, 30, 40; underfill ratio = 1.00%, Apex trigger(s) = 5–30, dynamic exclusion(s) = 15 s.

### LC-MS data analysis

RAW files were converted to mzML files using msconvert from ProteoWizard, using vendor centroiding on all scans, and analyzed using MAVEN2 software (Seitzer et al., 2022) against *in-house* libraries.

## Acknowledgements

We thank members of the Vander Heiden laboratory for helpful discussions and experimental advice. We acknowledge Paul-Jones Sali and Emily A. Dennstedt for mouse husbandry and technical assistance. E.C.L. was supported by the Damon Runyon Cancer Research Foundation (DRG-2299-17), the NIH (R00CA255928), and the Van Andel Institute Metabolism and Nutrition (MeNu) Program. A.N.L. was a Robert Black Fellow of the Damon Runyon Cancer Research Foundation (DRG-2241-15) and was supported by a NIH Pathway to Independence Award (K99CA234221). L.V.D. was supported by an NIH Ruth Kirschstein Fellowship (F32CA210421). M.G.V.H. acknowledges support from the Emerald Foundation, the Lustgarten Foundation, a Faculty Scholar grant from the Howard Hughes Medical Institute, Stand Up To Cancer, the MIT Center for Precision Cancer Medicine, the Ludwig Center at MIT, the NIH (R35CA242379, R01CA168653, R01CA201276, P30CA14051), and Calico Life Sciences.

## References

Altea-Manzano P, Doglioni G, Liu Y, Cuadros AM, Nolan E, Fernández-García J, Wu Q, Planque M, Laue KJ, Cidre-Aranaz F, Liu X-Z, Marin-Bejar O, Van Elsen J, Vermeire I, Broekaert D, Demeyer S, Spotbeen X, Idkowiak J, Montagne A, Demicco M, Alkan HF, Rabas N, Riera-Domingo C, Richard F, Geukens T, De Schepper M, Leduc S, Hatse S, Lambrechts Y, Kay EJ, Lilla S, Alekseenko A, Geldhof V, Boeckx B, de la Calle Arregui C, Floris G, Swinnen JV, Marine J-C, Lambrechts D, Pelechano V, Mazzone M, Zanivan S, Cools J, Wildiers H, Baud V, Grünewald TGP, Ben-David U, Desmedt C, Malanchi I, Fendt S-M. 2023. A palmitate-rich metastatic niche enables metastasis growth via p65 acetylation resulting in pro-metastatic NF-κB signaling. Nat Cancer. doi:10.1038/s43018-023-00513-2

Amorim JA, Coppotelli G, Rolo AP, Palmeira CM, Ross JM, Sinclair DA. 2022. Mitochondrial and metabolic dysfunction in ageing and age-related diseases. Nat Rev Endocrinol 18:243–258. doi:10.1038/s41574-021-00626-7

Anderson RM, Weindruch R. 2010. Metabolic reprogramming, caloric restriction and aging. Trends Endocrinol Metab 21:134–41. doi:10.1016/j.tem.2009.11.005

Antoniewicz MR, Kelleher JK, Stephanopoulos G. 2011. Measuring deuterium enrichment of glucose hydrogen atoms by gas chromatography/mass spectrometry. Anal Chem 83:3211–6. doi:10.1021/ac200012p

Bao XR, Ong S-E, Goldberger O, Peng J, Sharma R, Thompson DA, Vafai SB, Cox AG, Marutani E, Ichinose F, Goessling W, Regev A, Carr SA, Clish CB, Mootha VK. 2016. Mitochondrial dysfunction remodels one-carbon metabolism in human cells. Elife 5:e10575. doi:10.7554/eLife.10575

Bartman CR, Faubert B, Rabinowitz JD, DeBerardinis RJ. 2023. Metabolic pathway analysis using stable isotopes in patients with cancer. Nat Rev Cancer. doi:10.1038/s41568-023-00632-z

Bartman CR, TeSlaa T, Rabinowitz JD. 2021. Quantitative flux analysis in mammals. Nat Metab 3:896–908. doi:10.1038/s42255-021-00419-2

Betarbet R, Sherer TB, MacKenzie G, Garcia-Osuna M, Panov AV, Greenamyre JT. 2000. Chronic systemic pesticide exposure reproduces features of Parkinson’s disease. Nat Neurosci 3:1301–6. doi:10.1038/81834

Birsoy K, Wang T, Chen WW, Freinkman E, Abu-Remaileh M, Sabatini DM. 2015. An Essential Role of the Mitochondrial Electron Transport Chain in Cell Proliferation Is to Enable Aspartate Synthesis. Cell 162:540–51. doi:10.1016/j.cell.2015.07.016

Buescher JM, Antoniewicz MR, Boros LG, Burgess SC, Brunengraber H, Clish CB, DeBerardinis RJ, Feron O, Frezza C, Ghesquiere B, Gottlieb E, Hiller K, Jones RG, Kamphorst JJ, Kibbey RG, Kimmelman AC, Locasale JW, Lunt SY, Maddocks OD, Malloy C, Metallo CM, Meuillet EJ, Munger J, Nöh K, Rabinowitz JD, Ralser M, Sauer U, Stephanopoulos G, St-Pierre J, Tennant DA, Wittmann C, Vander Heiden MG, Vazquez A, Vousden K, Young JD, Zamboni N, Fendt SM. 2015. A roadmap for interpreting (13)C metabolite labeling patterns from cells. Curr Opin Biotechnol 34:189–201. doi:10.1016/j.copbio.2015.02.003

Chaurasia B, Summers SA. 2015. Ceramides - Lipotoxic Inducers of Metabolic Disorders. Trends Endocrinol Metab 26:538–550. doi:10.1016/j.tem.2015.07.006

Couttas TA, Kain N, Tran C, Chatterton Z, Kwok JB, Don AS. 2018. Age-Dependent Changes to Sphingolipid Balance in the Human Hippocampus are Gender-Specific and May Sensitize to Neurodegeneration. J Alzheimers Dis 63:503–514. doi:10.3233/JAD-171054

Davidson SM, Papagiannakopoulos T, Olenchock BA, Heyman JE, Keibler MA, Luengo A, Bauer MR, Jha AK, O’Brien JP, Pierce KA, Gui DY, Sullivan LB, Wasylenko TM, Subbaraj L, Chin CR, Stephanopolous G, Mott BT, Jacks T, Clish CB, Vander Heiden MG. 2016. Environment Impacts the Metabolic Dependencies of Ras-Driven Non-Small Cell Lung Cancer. Cell Metab 23:517–28. doi:10.1016/j.cmet.2016.01.007

Diehl FF, Lewis CA, Fiske BP, Vander Heiden MG. 2019. Cellular redox state constrains serine synthesis and nucleotide production to impact cell proliferation. Nat Metab 1:861–867. doi:10.1038/s42255-019-0108-x

Duan L, Cooper DE, Scheidemantle G, Locasale JW, Kirsch DG, Liu X. 2022. 13C tracer analysis suggests extensive recycling of endogenous CO2 in vivo. Cancer Metab 10:11. doi:10.1186/s40170-022-00287-8

Ehrhardt N, Cui J, Dagdeviren S, Saengnipanthkul S, Goodridge HS, Kim JK, Lantier L, Guo X, Chen Y-DI, Raffel LJ, Buchanan TA, Hsueh WA, Rotter JI, Goodarzi MO, Péterfy M. 2019. Adiposity-Independent Effects of Aging on Insulin Sensitivity and Clearance in Mice and Humans: Insulin Sensitivity in Aging. Obesity 27:434–443. doi:10.1002/oby.22418

Elia I, Rossi M, Stegen S, Broekaert D, Doglioni G, van Gorsel M, Boon R, Escalona-Noguero C, Torrekens S, Verfaillie C, Verbeken E, Carmeliet G, Fendt SM. 2019. Breast cancer cells rely on environmental pyruvate to shape the metastatic niche. Nature 568:117–121. doi:10.1038/s41586-019-0977-x

Fang EF, Kassahun H, Croteau DL, Scheibye-Knudsen M, Marosi K, Lu H, Shamanna RA, Kalyanasundaram S, Bollineni RC, Wilson MA, Iser WB, Wollman BN, Morevati M, Li J, Kerr JS, Lu Q, Waltz TB, Tian J, Sinclair DA, Mattson MP, Nilsen H, Bohr VA. 2016. NAD(+) Replenishment Improves Lifespan and Healthspan in Ataxia Telangiectasia Models via Mitophagy and DNA Repair. Cell Metab 24:566–581. doi:10.1016/j.cmet.2016.09.004

Faubert B, Li KY, Cai L, Hensley CT, Kim J, Zacharias LG, Yang C, Do QN, Doucette S, Burguete D, Li H, Huet G, Yuan Q, Wigal T, Butt Y, Ni M, Torrealba J, Oliver D, Lenkinski RE, Malloy CR, Wachsmann JW, Young JD, Kernstine K, DeBerardinis RJ. 2017. Lactate Metabolism in Human Lung Tumors. Cell 171:358–371.e9. doi:10.1016/j.cell.2017.09.019

Faubert B, Tasdogan A, Morrison SJ, Mathews TP, DeBerardinis RJ. 2021. Stable isotope tracing to assess tumor metabolism in vivo. Nat Protoc 16:5123–5145. doi:10.1038/s41596-021-00605-2

Gomes AP, Price NL, Ling AJ, Moslehi JJ, Montgomery MK, Rajman L, White JP, Teodoro JS, Wrann CD, Hubbard BP, Mercken EM, Palmeira CM, de Cabo R, Rolo AP, Turner N, Bell EL, Sinclair DA. 2013. Declining NAD(+) induces a pseudohypoxic state disrupting nuclear-mitochondrial communication during aging. Cell 155:1624–38. doi:10.1016/j.cell.2013.11.037

Goyal MS, Vlassenko AG, Blazey TM, Su Y, Couture LE, Durbin TJ, Bateman RJ, Benzinger TL, Morris JC, Raichle ME. 2017. Loss of Brain Aerobic Glycolysis in Normal Human Aging. Cell Metab 26:353–360.e3. doi:10.1016/j.cmet.2017.07.010

Green DR, Galluzzi L, Kroemer G. 2011. Mitochondria and the Autophagy– Inflammation–Cell Death Axis in Organismal Aging. Science 333:1109–1112. doi:10.1126/science.1201940

Heinrich P, Kohler C, Ellmann L, Kuerner P, Spang R, Oefner PJ, Dettmer K. 2018. Correcting for natural isotope abundance and tracer impurity in MS-, MS/MS- and high-resolution-multiple-tracer-data from stable isotope labeling experiments with IsoCorrectoR. Sci Rep 8:17910. doi:10.1038/s41598-018-36293-4

Hensley CT, Faubert B, Yuan Q, Lev-Cohain N, Jin E, Kim J, Jiang L, Ko B, Skelton R, Loudat L, Wodzak M, Klimko C, McMillan E, Butt Y, Ni M, Oliver D, Torrealba J, Malloy CR, Kernstine K, Lenkinski RE, DeBerardinis RJ. 2016. Metabolic Heterogeneity in Human Lung Tumors. Cell 164:681–94. doi:10.1016/j.cell.2015.12.034

Hertel J, Friedrich N, Wittfeld K, Pietzner M, Budde K, Van der Auwera S, Lohmann T, Teumer A, Völzke H, Nauck M, Grabe HJ. 2016. Measuring Biological Age via Metabonomics: The Metabolic Age Score. J Proteome Res 15:400–10. doi:10.1021/acs.jproteome.5b00561

Houtkooper RH, Argmann C, Houten SM, Cantó C, Jeninga EH, Andreux PA, Thomas C, Doenlen R, Schoonjans K, Auwerx J. 2011. The metabolic footprint of aging in mice. Sci Rep 1:134. doi:10.1038/srep00134

Houtkooper RH, Pirinen E, Auwerx J. 2012. Sirtuins as regulators of metabolism and healthspan. Nat Rev Mol Cell Biol 13:225–238. doi:10.1038/nrm3293

Hoxhaj G, Manning BD. 2020. The PI3K-AKT network at the interface of oncogenic signalling and cancer metabolism. Nat Rev Cancer 20:74–88. doi:10.1038/s41568-019-0216-7

Hui S, Cowan AJ, Zeng X, Yang L, TeSlaa T, Li X, Bartman C, Zhang Z, Jang C, Wang L, Lu W, Rojas J, Baur J, Rabinowitz JD. 2020. Quantitative Fluxomics of Circulating Metabolites. Cell Metab. doi:10.1016/j.cmet.2020.07.013

Hui S, Ghergurovich JM, Morscher RJ, Jang C, Teng X, Lu W, Esparza LA, Reya T, Le Z, Yanxiang Guo J, White E, Rabinowitz JD. 2017. Glucose feeds the TCA cycle via circulating lactate. Nature 551:115–118. doi:10.1038/nature24057

Hulbert AJ, Pamplona R, Buffenstein R, Buttemer WA. 2007. Life and death: metabolic rate, membrane composition, and life span of animals. Physiol Rev 87:1175–213. doi:10.1152/physrev.00047.2006

Jang C, Hui S, Lu W, Cowan AJ, Morscher RJ, Lee G, Liu W, Tesz GJ, Birnbaum MJ, Rabinowitz JD. 2018. The Small Intestine Converts Dietary Fructose into Glucose and Organic Acids. Cell Metab 27:351–361.e3. doi:10.1016/j.cmet.2017.12.016

Jang C, Hui S, Zeng X, Cowan AJ, Wang L, Chen L, Morscher RJ, Reyes J, Frezza C, Hwang HY, Imai A, Saito Y, Okamoto K, Vaspoli C, Kasprenski L, Zsido GA, Gorman JH, Gorman RC, Rabinowitz JD. 2019. Metabolite Exchange between Mammalian Organs Quantified in Pigs. Cell Metab 30:594–606.e3. doi:10.1016/j.cmet.2019.06.002

Jové M, Naudí A, Ramírez-Núñez O, Portero-Otín M, Selman C, Withers DJ, Pamplona R. 2014. Caloric restriction reveals a metabolomic and lipidomic signature in liver of male mice. Aging Cell 13:828–37. doi:10.1111/acel.12241

Kim W, Deik A, Gonzalez C, Gonzalez ME, Fu F, Ferrari M, Churchhouse CL, Florez JC, Jacobs SBR, Clish CB, Rhee EP. 2019. Polyunsaturated Fatty Acid Desaturation Is a Mechanism for Glycolytic NAD^+^ Recycling. Cell Metab. doi:10.1016/j.cmet.2018.12.023

Kishimoto Y, Radin NS. 1959. Composition of cerebroside acids as a function of age. Journal of Lipid Research 1:79–82. doi:10.1016/S0022-2275(20)39096-9

Lau AN, Li Z, Danai LV, Westermark AM, Darnell AM, Ferreira R, Gocheva V, Sivanand S, Lien EC, Sapp KM, Mayers JR, Biffi G, Chin CR, Davidson SM, Tuveson DA, Jacks T, Matheson NJ, Yilmaz O, Vander Heiden MG. 2020. Dissecting cell-type-specific metabolism in pancreatic ductal adenocarcinoma. Elife 9:e56782. doi:10.7554/eLife.56782

Laye MJ, Tran V, Jones DP, Kapahi P, Promislow DE. 2015. The effects of age and dietary restriction on the tissue-specific metabolome of Drosophila. Aging Cell 14:797–808. doi:10.1111/acel.12358

Li Z, Ji BW, Dixit PD, Tchourine K, Lien EC, Hosios AM, Abbott KL, Rutter JC, Westermark AM, Gorodetsky EF, Sullivan LB, Vander Heiden MG, Vitkup D. 2022. Cancer cells depend on environmental lipids for proliferation when electron acceptors are limited. Nat Metab 4:711–723. doi:10.1038/s42255-022-00588-8

Lien EC, Westermark AM, Zhang Y, Yuan C, Li Z, Lau AN, Sapp KM, Wolpin BM, Vander Heiden MG. 2021. Low glycaemic diets alter lipid metabolism to influence tumour growth. Nature 599:302–307. doi:10.1038/s41586-021-04049-2

López-Otín C, Blasco MA, Partridge L, Serrano M, Kroemer G. 2013. The hallmarks of aging. Cell 153:1194–217. doi:10.1016/j.cell.2013.05.039

López-Otín C, Galluzzi L, Freije JM, Madeo F, Kroemer G. 2016. Metabolic Control of Longevity. Cell 166:802–21. doi:10.1016/j.cell.2016.07.031

Luengo A, Li Z, Gui DY, Sullivan LB, Zagorulya M, Do BT, Ferreira R, Naamati A, Ali A, Lewis CA, Thomas CJ, Spranger S, Matheson NJ, Vander Heiden MG. 2021. Increased demand for NAD^+^ relative to ATP drives aerobic glycolysis. Mol Cell 81:691–707.e6. doi:10.1016/j.molcel.2020.12.012

Matyash V, Liebisch G, Kurzchalia TV, Shevchenko A, Schwudke D. 2008. Lipid extraction by methyl-tert-butyl ether for high-throughput lipidomics. J Lipid Res 49:1137–1146. doi:10.1194/jlr.D700041-JLR200

McReynolds MR, Chellappa K, Baur JA. 2020. Age-related NAD+ decline. Exp Gerontol 134:110888. doi:10.1016/j.exger.2020.110888

McReynolds MR, Chellappa K, Chiles E, Jankowski C, Shen Y, Chen L, Descamps HC, Mukherjee S, Bhat YR, Lingala SR, Chu Q, Botolin P, Hayat F, Doke T, Susztak K, Thaiss CA, Lu W, Migaud ME, Su X, Rabinowitz JD, Baur JA. 2021. NAD^+^ flux is maintained in aged mice despite lower tissue concentrations. Cell Syst S2405-4712(21)00338–0. doi:10.1016/j.cels.2021.09.001

Miller KN, Burhans MS, Clark JP, Howell PR, Polewski MA, DeMuth TM, Eliceiri KW, Lindstrom MJ, Ntambi JM, Anderson RM. 2017. Aging and caloric restriction impact adipose tissue, adiponectin, and circulating lipids. Aging Cell 16:497–507. doi:10.1111/acel.12575

Mills KF, Yoshida S, Stein LR, Grozio A, Kubota S, Sasaki Y, Redpath P, Migaud ME, Apte RS, Uchida K, Yoshino J, Imai SI. 2016. Long-Term Administration of Nicotinamide Mononucleotide Mitigates Age-Associated Physiological Decline in Mice. Cell Metab 24:795–806. doi:10.1016/j.cmet.2016.09.013

Naudí A, Jové M, Ayala V, Portero-Otín M, Barja G, Pamplona R. 2013. Membrane lipid unsaturation as physiological adaptation to animal longevity. Front Physiol 4:372. doi:10.3389/fphys.2013.00372

Reynolds TH, Dalton A, Calzini L, Tuluca A, Hoyte D, Ives SJ. 2019. The impact of age and sex on body composition and glucose sensitivity in C57BL/6J mice. Physiol Rep 7:e13995. doi:10.14814/phy2.13995

Ross JM, Öberg J, Brené S, Coppotelli G, Terzioglu M, Pernold K, Goiny M, Sitnikov R, Kehr J, Trifunovic A, Larsson NG, Hoffer BJ, Olson L. 2010. High brain lactate is a hallmark of aging and caused by a shift in the lactate dehydrogenase A/B ratio. Proc Natl Acad Sci U S A 107:20087–92. doi:10.1073/pnas.1008189107

Seitzer P, Bennett B, Melamud E. 2022. MAVEN2: An Updated Open-Source Mass Spectrometry Exploration Platform. Metabolites 12:684. doi:10.3390/metabo12080684

Sellers K, Fox MP, Bousamra M, Slone SP, Higashi RM, Miller DM, Wang Y, Yan J, Yuneva MO, Deshpande R, Lane AN, Fan TW. 2015. Pyruvate carboxylase is critical for non-small-cell lung cancer proliferation. J Clin Invest 125:687–98. doi:10.1172/JCI72873

Sullivan LB, Gui DY, Hosios AM, Bush LN, Freinkman E, Vander Heiden MG. 2015. Supporting Aspartate Biosynthesis Is an Essential Function of Respiration in Proliferating Cells. Cell 162:552–63. doi:10.1016/j.cell.2015.07.017

Titov DV, Cracan V, Goodman RP, Peng J, Grabarek Z, Mootha VK. 2016. Complementation of mitochondrial electron transport chain by manipulation of the NAD+/NADH ratio. Science 352:231–5. doi:10.1126/science.aad4017

Vander Heiden MG, DeBerardinis RJ. 2017. Understanding the Intersections between Metabolism and Cancer Biology. Cell 168:657–669. doi:10.1016/j.cell.2016.12.039

Verdin E. 2015. NAD^+^ in aging, metabolism, and neurodegeneration. Science 350:1208–13. doi:10.1126/science.aac4854

Vriens K, Christen S, Parik S, Broekaert D, Yoshinaga K, Talebi A, Dehairs J, Escalona-Noguero C, Schmieder R, Cornfield T, Charlton C, Romero-Pérez L, Rossi M, Rinaldi G, Orth MF, Boon R, Kerstens A, Kwan SY, Faubert B, Méndez-Lucas A, Kopitz CC, Chen T, Fernandez-Garcia J, Duarte JAG, Schmitz AA, Steigemann P, Najimi M, Hägebarth A, Van Ginderachter JA, Sokal E, Gotoh N, Wong KK, Verfaillie C, Derua R, Munck S, Yuneva M, Beretta L, DeBerardinis RJ, Swinnen JV, Hodson L, Cassiman D, Verslype C, Christian S, Grünewald S, Grünewald TGP, Fendt SM. 2019. Evidence for an alternative fatty acid desaturation pathway increasing cancer plasticity. Nature 566:403–406. doi:10.1038/s41586-019-0904-1

Walters RO, Arias E, Diaz A, Burgos ES, Guan F, Tiano S, Mao K, Green CL, Qiu Y, Shah H, Wang D, Hudgins AD, Tabrizian T, Tosti V, Shechter D, Fontana L, Kurland IJ, Barzilai N, Cuervo AM, Promislow DEL, Huffman DM. 2018. Sarcosine Is Uniquely Modulated by Aging and Dietary Restriction in Rodents and Humans. Cell Rep 25:663–676.e6. doi:10.1016/j.celrep.2018.09.065

Wang B, Tontonoz P. 2019. Phospholipid Remodeling in Physiology and Disease. Annu Rev Physiol 81:165–188. doi:10.1146/annurev-physiol-020518-114444

Weetman J, Wong MB, Sharry S, Rcom-H’cheo-Gauthier A, Gai WP, Meedeniya A, Pountney DL. 2013. Increased SUMO-1 expression in the unilateral rotenone-lesioned mouse model of Parkinson’s disease. Neurosci Lett 544:119–24. doi:10.1016/j.neulet.2013.03.057

Yuan R, Tsaih S-W, Petkova SB, De Evsikova CM, Xing S, Marion MA, Bogue MA, Mills KD, Peters LL, Bult CJ, Rosen CJ, Sundberg JP, Harrison DE, Churchill GA, Paigen B. 2009. Aging in inbred strains of mice: study design and interim report on median lifespans and circulating IGF1 levels: Median lifespans and IGF1 levels of 31 inbred strains. Aging Cell 8:277–287. doi:10.1111/j.1474-9726.2009.00478.x

Yuneva MO, Fan TW, Allen TD, Higashi RM, Ferraris DV, Tsukamoto T, Mates JM, Alonso FJ, Wang C, Seo Y, Chen X, Bishop JM. 2012. The metabolic profile of tumors depends on both the responsible genetic lesion and tissue type. Cell Metab 15:157–70. doi:10.1016/j.cmet.2011.12.015

