## Supplemental Figure Legends for "Effects of aging on glucose and lipid metabolism in mice"

**Figure S1. A 6-hour infusion is required to reach steady-state labeling in C57BL/6J mouse tissues.** C57BL/6J mice were infused with [U-$^{13}$C]-glucose at 0.4 mg/min for 0.5 h (n = 2), 2 h (n = 3), 4 h (n = 3), and 6 h (n = 2 for plasma, n = 3 for tissues). **A,** Plasma glucose enrichment. **B-D,** [M+3] fractional labeling of pyruvate, lactate, and alanine (**B**), [M+2] fractional labeling of the indicated TCA cycle metabolites (**C**), and fractional labeling of the indicated amino acids (**D**) in plasma, liver, gastrocnemius muscle, and brain tissues over time. Data are presented as mean ± SEM.

**Figure S2. [U-$^{13}$C]-glucose infusion rate differences can impact tissue metabolite labeling patterns in young versus old C57BL/6J mice.** 3 month old (n = 3) versus 24 month old (n = 6) C57BL/6J mice were infused with [U-$^{13}$C]-glucose at 30 mg/kg/min for 4 h. **A, B,** [M+3] fractional labeling of pyruvate (**A**) and lactate (**B**) in the indicated tissues. Data are presented as mean ± SEM. Comparisons were made using a two-tailed Student's t-test.

**Figure S3. Glucose contribution to glycolysis is robust in aging WSB/EiJ and DO mice.** 3 month old (n = 6) versus 24 month old (n = 5) WSB/EiJ mice were infused with [U-$^{13}$C]-glucose at 0.4 mg/min for 6 h. 3 month old (n = 6), 24 month old (n = 4), and 30 month old (n = 4) DO mice were infused with [U-$^{13}$C]-glucose at 0.4 mg/min for 6 h. **A-C,** [M+3] fractional labeling of pyruvate (**A**), lactate (**B**), and alanine (**C**) in the indicated tissues from WSB/EiJ mice. **D-F,** Relative levels of pyruvate (**D**), lactate (**E**), and alanine

(**F**) in the indicated tissues from WSB/EiJ mice. **G-I,** [M+3] fractional labeling of pyruvate (**G**), lactate (**H**), and alanine (**I**) in the indicated tissues from DO mice. **J-L,** Relative levels of pyruvate (**J**), lactate (**K**), and alanine (**L**) in the indicated tissues from DO mice. Data are presented as mean ± SEM. Comparisons were made using a two-tailed Student's t-test.

**Figure S4. Glucose contribution to the TCA cycle is robust in aging C57BL/6J mice.** 3 month old (n = 4) versus 24 month old (n = 4) C57BL/6J mice were infused with [U-$^{13}$C]-glucose at 0.4 mg/min for 6 h. **A-D,** Mass isotopomer distributions of α-ketoglutarate (αKG), fumarate, and aspartate in plasma (**A**), liver (**B**), gastrocnemius muscle (**C**), and brain (**D**) tissues. Data are presented as mean ± SEM.

**Figure S5. Glucose contribution to the TCA cycle is robust in aging WSB/EiJ mice.** 3 month old (n = 6) versus 24 month old (n = 5) WSB/EiJ mice were infused with [U-$^{13}$C]-glucose at 0.4 mg/min for 6 h. **A-D,** Relative levels of TCA cycle metabolites in plasma (**A**), liver (**B**), gastrocnemius muscle (**C**), and brain (**D**) tissues. **E-H,** Mass isotopomer distributions of the indicated TCA cycle metabolites in plasma (**E**), liver (**F**), gastrocnemius muscle (**G**), and brain (**H**) tissues. Data are presented as mean ± SEM. Comparisons were made using a two-tailed Student's t-test.

**Figure S6. Glucose contribution to the TCA cycle is robust in aging DO mice.** 3 month old (n = 6), 24 month old (n = 4), and 30 month old (n = 4) DO mice were infused with [U-$^{13}$C]-glucose at 0.4 mg/min for 6 h. **A-D,** Relative levels of TCA cycle metabolites in plasma (**A**), liver (**B**), gastrocnemius muscle (**C**), and brain (**D**) tissues. **E-H,** Mass isotopomer distributions of the indicated TCA cycle metabolites in plasma (**E**), liver (**F**), gastrocnemius muscle (**G**), and brain (**H**) tissues. Data are presented as mean ± SEM. Comparisons were made using a two-tailed Student's t-test.

**Figure S7. Glucose contribution to amino acid metabolism is robust in aging WSB/EiJ mice.** 3 month old (n = 6) versus 24 month old (n = 5) WSB/EiJ mice were infused with [U-$^{13}$C]-glucose at 0.4 mg/min for 6 h. **A,** Relative levels of the indicated amino acids in plasma, liver, gastrocnemius muscle, and brain tissues. **B-G,** Fractional labeling of [M+2] and [M+3] asparagine (**B**), [M+2] and [M+3] glutamine (**C**), [M+2] and [M+3] glutamate (**D**), [M+2] and [M+3] proline (**E**), [M+3] and [M+1] serine (**F**), and [M+2] glycine (**G**) in the indicated tissues. Data are presented as mean ± SEM. Comparisons were made using a two-tailed Student's t-test.

**Figure S8. Glucose contribution to amino acid metabolism is robust in aging DO mice.** 3 month old (n = 6), 24 month old (n = 4), and 30 month old (n = 4) DO mice were infused with [U-$^{13}$C]-glucose at 0.4 mg/min for 6 h. **A,** Relative levels of the indicated amino acids in plasma, liver, gastrocnemius muscle, and brain tissues. **B-G,** Fractional labeling of [M+2] and [M+3] asparagine (**B**), [M+2] and [M+3] glutamine (**C**), [M+2] and [M+3] glutamate (**D**), [M+2] and [M+3] proline (**E**), [M+3] and [M+1] serine (**F**), and [M+2] glycine (**G**) in the indicated tissues. Data are presented as mean ± SEM. Comparisons were made using a two-tailed Student's t-test.

**Figure S9. Polar metabolite profiling does not reveal strong age-dependent changes in metabolite levels.** Polar metabolite levels were measured by LC-MS in tissues from uninfused 3 month old (n = 9), 12 month old (n = 3), and 24 month old (n = 3) C57BL/6J mice. Data were analyzed using MetaboAnalyst. **A-C,** Heat maps of metabolite levels from liver (**A**), gastrocnemius muscle (**B**), and brain (**C**) tissues. **D-F,** Principal component analysis of metabolite levels from liver (**D**), gastrocnemius muscle (**E**), and brain (**F**) tissues.

**Figure S10. Fatty acid desaturation increases in tissues from aging WSB/EiJ mice. A-D,** Relative levels of the indicated fatty acids in plasma (**A**), liver (**B**), gastrocnemius muscle (**C**), and brain (**D**) tissues from young versus old WSB/EiJ mice. **E-H,** Relative fatty acid desaturation ratios in plasma (**E**), liver (**F**), gastrocnemius muscle (**G**), and brain (**H**) tissues from young versus old WSB/EiJ mice. 3 months n = 6, 24 months n = 5. Data are presented as mean ± SEM. Comparisons were made using a two-tailed Student's t-test.

**Figure S11. Fatty acid desaturation increases in tissues from aging DO mice. A-D,** Relative levels of the indicated fatty acids in plasma (**A**), liver (**B**), gastrocnemius muscle (**C**), and brain (**D**) tissues from young versus old DO mice. **E-H,** Relative fatty acid desaturation ratios in plasma (**E**), liver (**F**), gastrocnemius muscle (**G**), and brain (**H**) tissues from young versus old DO mice. 3 months n = 6, 24 months n = 4, 30 months n = 4. Data are presented as mean ± SEM. Comparisons were made using a two-tailed Student's t-test.

**Figure S12. Aging brain tissue exhibits changes in levels of d18:0-, d18:1-, and d18:2-containing sphingolipid species. A-C,** Heat maps of relative levels of d18:0-, d18:1-, and d18:2-containing sphingolipid species in brain tissues from young versus old C57BL/6J (**A**), WSB/EiJ (**B**), and DO (**C**) mice. C57BL/6J: 3 months n = 3, 24 months n = 3. WSB/EiJ: 3 months n = 6, 24 months n = 5. DO: 3 months n = 6, 24 months n = 4, 30 months n = 4.

**Figure S13. Aging brain tissue exhibits changes in levels of sphingolipid species that contain 2-hydroxylated fatty acids. A-C,** Heat maps of relative levels of sphingolipid species that contain 2-hydroxylated fatty acids in brain tissues from young versus old C57BL/6J (**A**), WSB/EiJ (**B**), and DO (**C**) mice. C57BL/6J: 3 months n = 3, 24 months n = 3. WSB/EiJ: 3 months n = 6, 24 months n = 5. DO: 3 months n = 6, 24 months n = 4, 30 months n = 4.
